# Do training with blurred images make convolutional neural networks closer to humans concerning object recognition performance and internal representations?

**DOI:** 10.1101/2022.06.13.496005

**Authors:** Sou Yoshihara, Taiki Fukiage, Shin’ya Nishida

## Abstract

It is suggested that experiences of perceiving blurry images in addition to sharp images contribute to the development of robust human visual processing. To computationally investigate the effect of exposure to blurry images, we trained Convolutional Neural Networks (CNNs) on ImageNet object recognition with a variety of combinations of sharp and blurry images. In agreement with related studies, mixed training on sharp and blurred images (B+S) makes the CNNs close to humans with respect to robust object recognition against a change in image blur. B+S training also reduces the texture bias of CNN in recognition of shape-texture-cue-conflict images, but the effect is not strong enough to achieve a strong shape bias comparable to what humans show. Other tests also suggest that B+S training is not sufficient to produce robust human-like object recognition based on global configurational features. We also show using representational similarity analysis and zero-shot transfer learning that B+S-Net does not acquire blur-robust object recognition through separate specialized sub-networks, each for sharp and blurry images, but through a single network analyzing common image features. However, blur training alone does not automatically create a mechanism like the human brain where subband information is integrated into a common representation. Our analyses suggest that experience with blurred images helps the human brain develop neural networks that robustly recognize the surrounding world, but it is not powerful enough to fill a large gap between humans and CNNs.

## 1 Introduction

Human visual acuity, evaluated in terms of the minimum angle of resolution or the highest discernable spatial frequency, is affected by a variety of processes including eye optics, retinal sensor sampling, and the subsequent neural signal processing. In daily visual experiences, visual acuity changes depending on, for example, how much the current focal length of the eye agrees with the distance to the target object, or whether the target is sensed at the fovea where image sampling is dense or at far-peripheral vision where sparsely image sampling is followed by spatially pooling. Visual acuity also changes with developmental stages. Infants who were born with low visual acuity gradually acquire near adult-level acuity within the first few years of their life [Dobson and Teller, 1978, Banks and Salapatek, 1981]. Considering that the loss of visual acuity can be approximated by blurring the image by low-pass (high-cut) filtering, one can say that most humans have rich experience seeing blurred visual images in addition to sharp ones.

It has been suggested that the experience of blurred visual images might be functionally beneficial, enabling the visual system to use global configural structures in image recognition [Grand et al., 2001, Le Grand et al., 2004, Vogelsang et al., 2018]. Several recent studies use machine learning of artificial neural networks to computationally test this hypothesis [Vogelsang et al., 2018, Katzhendler and Weinshall, 2019, Avberšek et al., 2021, Jang and Tong, 2021]. Vogelsang trained the convolutional network (CNN) to recognize human faces. To simulates a gradual improvement of visual acuity during the initial stage of life, they changed the training images from blurred to sharp ones during training (B2S) and found that the network achieves robust face recognition for a wide range of image blur, as humans do. In contrast, the network can only recognize sharp images when trained on sharp images. The network can only recognize blurred images when trained using blurred images or sequentially trained on images changing from sharp to blurred images. Jang and Tong found that the effect of the B2S training is task-specific. It leads to blur-robust recognition for face recognition as Vogelsang et al. reported, but not for object recognition. Avberšek et al. neither obtained blur-robust object recognition by the B2S training. However, object recognition achieves blur-robustness when blurred and sharp images are always mixed during training (B+S).

With a similar research motivation in mind, we examined the effects of blur training on object recognition by CNNs. We investigated which types of blur training make the CNNs sensitive to coarse-scale global features as well as fine-scale local features, and make them close to the human object recognition system. We evaluated the object-recognition performance of the blur-trained CNNs not only using low-pass filtered test images, but also for other types of images including bandpass filtered images and shape-texture-cue-conflict images [Geirhos et al., 2019] to see whether blur training affects global configurational processing in general. In agreement with the previous reports [Avberšek et al., 2021, Jang and Tong, 2021], our results show that mixed training on sharp and blurred images (B+S) is more effective in making the CNNs robust against a change in image blur in object recognition in comparison with the training simulating the human development by gradually improving image sharpness during training (B2S). The B+S training however is not sufficient to produce robust human-like object recognition based on global configurational features. For example, it reduces the texture bias of CNN for shape-texture-cue-conflict images, but the effect is too small to achieve a strong shape bias comparable to what humans show.

In the latter half of this report, using correlation analyses of internal representations and zero-shot transfer learning, we examined how B+S training makes CNN robust against a range of image blur. Our results suggest that the blur robustness of B+S-Net is contributed by initial low-pass filtering, but only partially. Representational similarities in the intermediate layers suggest that B+S-Net processes sharp and blurry images not through separate specialized sub-networks, but through a common mechanism. Furthermore, we found that B+S training for other object labels transfers to another label trained only with blurred or sharp images, which suggests that B+S training lets the network learn general blur-robust features. However, blur training alone does not automatically create a mechanism like the human brain where subband information is integrated into a common representation.

Overall, our results suggest that experience with blurred images helps the human brain develop neural networks that robustly recognize the surrounding world, but its contribution to reducing a large gap between humans and CNNs is limited.

## 2 Methods

We investigated the performance of several training methods with a mixture of blurred images. In the experiments, we mainly used 16-class-ImageNet[Geirhos et al., 2018] as a dataset, and the analysis is based on 16-class-AlexNet, with 16 final layer units. However, we also ran some of the experiments using a 1000-class-ImageNet and tested other network architectures to ensure the generalizability of our results. A list of the networks compared in this study is summarized in Table 1. We trained all the models from scratch except for SIN-trained-Net[Geirhos et al., 2019], for which we used the pre-trained model provided by the authors. We did not fine-tune any of the models for the test tasks. Further, we collected human behavior data via Amazon Mechanical Turk (AMT) to compare human performances with those of our blur-trained models. Below, we describe the details about the dataset, model architecture, training strategies, and human behavior study.

**Table 1:** CNN models evaluated

| Model name | Architecture | Number of units in final layer | Training dataset | Pre-training | Fine-tuning |
| --- | --- | --- | --- | --- | --- |
| 16-class-AlexNet | AlexNet | 16 | 16-class-ImageNet[Geirhos et al., 2018] | none | none |
| 1000-class-AlexNet | AlexNet | 1000 | ImageNet(ILSVRC2012) | none | none |
| VOneNet | VOneBlock[Dapello et al., 2020] + AlexNet | 16 | 16-class-ImageNet[Geirhos et al., 2018] | none | none |
| 16-class-VGG16 | VGG16[Simonyan and Zisserman, 2015] | 16 | 16-class-ImageNet[Geirhos et al., 2018] | none | none |
| 16-class-ResNet50 | ResNet50[He et al., 2016] | 16 | 16-class-ImageNet[Geirhos et al., 2018] | none | none |
| SIN-trained-Net[Geirhos et al., 2019] | AlexNet | 1000 | SIN-ImageNet[Geirhos et al., 2019] | Yes | No |

### 2.1 dataset: 16-class-ImageNet

In order to facilitate comparison with experimental data on humans, we used the 16-class-ImageNet dataset. This dataset was created by Geirhos et al.[Geirhos et al., 2018], who grouped ImageNet classes into higher classes such as “dog” and “clock” and selected 16 of them. There are 40,517 training images and 1,600 test images. The image size is 224 × 224 × 3 (height, width, colour). The performance of the model trained on the regular 1000-class-ImageNet is also investigated in a later section (Section 3.5.1).

16-class name

airplane, bear, bicycle, bird, boat, bottle, car, cat, chair, clock, dog, elephant, keyboard, knife, oven, truck

### 2.2 CNN model: 16-class-AlexNet

We chose AlexNet[Krizhevsky et al., 2012] as the CNN model for our main analysis. We used a model architecture provided in a popular deep learning framework, Pytorch, and trained the model from scratch. To match the number of classes in the 16-class-ImageNet, we changed the number of output in the final layer from 1000 to 16.

We chose AlexNet because of its similarity to the hierarchical information processing of the human visual cortex. For example, the visualization of filters in the first layer of AlexNet trained with ImageNet shows the formation of various Gabor-like filters with different orientations and scales [Krizhevsky et al., 2012]. The Gabor functions are known to be good approximations of the spatial properties of V1 simple cell receptive fields [Jones and Palmer, 1987]. (In Section 3.5.2, we also analyze a model that explicitly incorporated the Gabor filters as the initial layer of AlexNet using VOneBlock proposed by [Dapello et al., 2020].) A study of the brain hierarchy (BH) score, which takes into account the hierarchical similarity between the deep neural network (DNN) and the brain, shows that AlexNet has a high BH score [Nonaka et al., 2020]. The information representation in the convolutional layer of AlexNet corresponds to the lower visual cortex of the brain, while the fully connected layer corresponds to the higher visual cortex of the brain. In addition, AlexNet is an easy model to interpret in that it has a small number of layers and does not contain complex operations such as Skip Connection.

### 2.3 Training with blurred images: Blur-training

In this experiment, in addition to the regular training, we trained CNNs with blurred images according to three different strategies (Figure 2). We applied Gaussian kernel convolution to blur images. The blur size was manipulated by changing the standard deviation (*σ*) of the Gaussian kernel as shown in Fig. 1. The spatial extent of the Gaussian kernel (*k*) was determined depending on *σ* as follows: *k* = *Round*(8*σ* + 1). ^1^. When *k* was an even number, one was added to make it an odd number.

**Figure 1:**
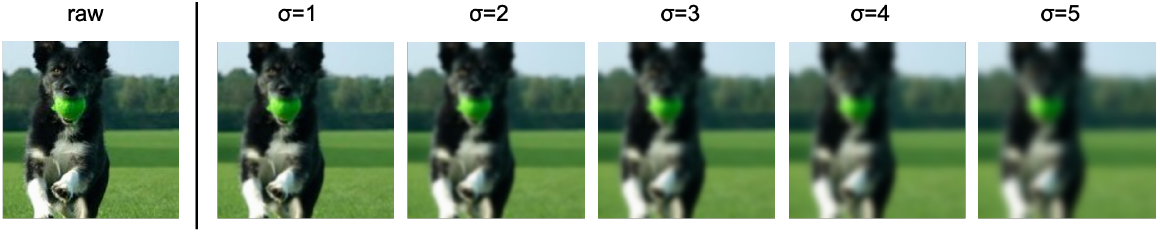
Image blur with Gaussian function

**Figure 2:**
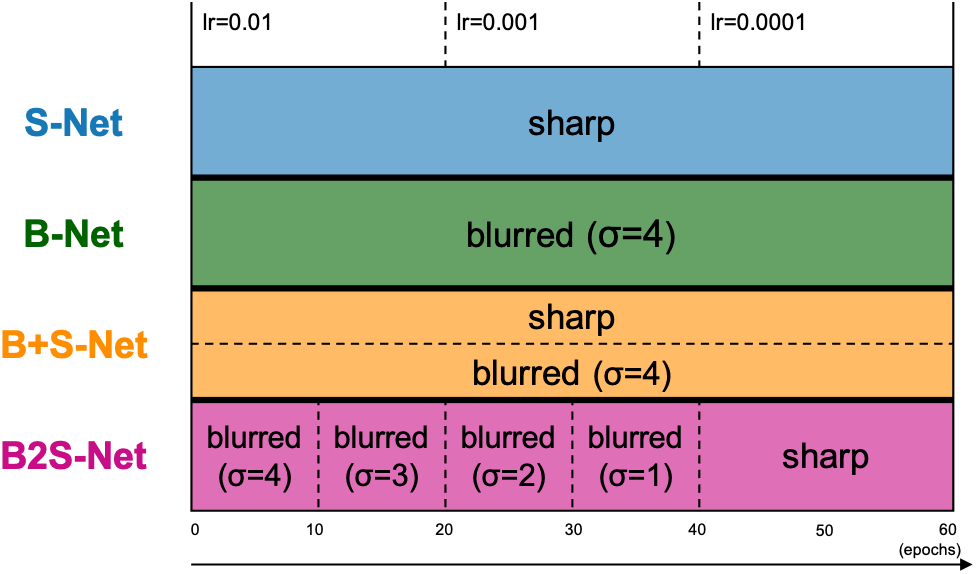
training method

In the following, we refer to the trained models as S-Net, B-Net, B+S-Net, and B2S-Net, respectively, according to the different image blurring strategies. Unless otherwise stated, the architecture of each model is 16-class-AlexNet. We trained all the models for 60 epochs (the number of training cycle through the full training dataset), with a batch size of 64. The optimizer was stochastic gradient descent (SGD) with momentum=0.9 and weight decay=0.0005. The initial learning rate (lr) was set to 0.01 and decreased by a factor of ten at every 20 epochs as shown in Figure 2. The number of training images was 40,517, the same for all models, and we applied random cropping and random horizontal flipping to all training images. The image size was 224 × 224 × 3 (height, width, colour). We used PyTorch (version 1.2.0) and one of two GPU machines to train each model. The GPU environments were Quadro RTX 8000 (CUDA Version: 10.2) and GeForce RTX 2080 (CUDA Version: 10.2).

#### S-Net

S-Net is a model trained on sharp (original, unblurred) images.

#### B-Net

In the training of B-Net, all the training images were blurred throughout the entire training period. We mainly discuss the performance of the model trained with the fixed blur size of *σ* = 4px.

#### B+S-Net

In the training of B+S-Net, we blurred half of the samples randomly picked in each batch of training images throughout the entire training period. We mainly discuss the performance of the model trained with the fixed blur size of *σ* = 4px. The performance of B+S-Net trained with randomly varied *σ* is presented in Section 3.4.

#### B2S-Net

In the training of B2S-Net, the training images were progressively made sharper from a strongly blurred to the original, non-blurred image. Specifically, we started with a Gaussian kernel of *σ* = 4px and decreased *σ* by one every ten epochs so that only sharp images without any blur were fed into the model in the last 20 epochs. This training method is intended to simulate human visual development and to confirm the effectiveness of starting training with blurred images, as claimed by Vogelsang et al.[Vogelsang et al., 2018].

### 2.4 Human image classification task

We asked participants to perform an image classification task on Amazon Mechanical Turk (AMT) to investigate the difference between the models trained in this study and human image recognition capabilities.

For stimuli of the classification task, we used the same 16-class-ImageNet test set that we used for evaluating CNN models (1600 images, 100 images per class). In addition to the original test images, we tested the low-pass and band-pass version of the 16-class-ImageNet test images for stimuli. The low-pass images were created by applying Gaussian kernel convolution while manipulating the standard deviation of the Gaussian kernel (*σ*) in the same manner as when we blurred the training images for CNN models. The band-pass images were created by taking the difference between two low-pass images obtained by blurring the same image with different *σ* (Figure 5). In total, there were six conditions as summarized below.

The six test conditions used in the human image classification task

Original image, low-pass image *σ* = 4*px*, low-pass image *σ* = 8*px*, low-pass image *σ* = 16*px*, band-pass image *σ*1*px* − *σ*2*px*, band-pass image *σ*4*px* − *σ*8*px*.

For each task, one of the stimuli was presented and participants chose the category of an object in the image from 16 options. We had 170 subjects solve the tasks and obtained 6,817 categorization data (Original image: 1108, low-pass image *σ* = 4*px*: 1103, low-pass image *σ* = 8*px*: 1124, low-pass image *σ* = 16*px*: 1134, band-pass image *σ*1*px* − *σ*2*px*: 1188, band-pass image *σ*4*px* − *σ*8*px*: 1160). Participants could complete the task for an arbitrary number of images. The consent form for the experiment was created using a Google Form, and a link to it was placed on the AMT task page. Each participant was asked to read the linked consent form and fill in the necessary information to give his or her consent. Experimental procedures were approved by the Research Ethics Committee at Graduate School of Informatics, Kyoto University, and were conducted in accordance with the Declaration of Helsinki.

## 3 Recognition performance comparison across different networks

We measure the model’s performance on various test images and compare it to human performance to investigate what visual functions are acquired via blur training. Firstly, we examined the classification accuracy for low spatial frequency images and analyzed whether the models could recognize coarse-scale information. Then, we examined whether the robustness to blurry images acquired through blur training could generalize to other types of robustness measured by using band-pass filtered images, images with manipulated spatial configuration of local elements, and the shape-texture cue conflict images.

### 3.1 Recognition performance for low-pass images

In this section, we compare the image recognition performance from low spatial frequency features. Since the low frequency information can capture global image features to some extent, the results of this task are expected to indicate, at least partially, whether the model recognizes global information or not. For this purpose, we examined the percentage of correct classification for each model when the test image was blurred at different intensities. The test images are the test set of the 16-class-ImageNet containing 1600 images. We also collected human classification task data using the same test images. The details are described in Section 2.4.

The results of the above experiments are shown in Figure 4. First, S-Net trained only on standard clear images shows a sharp drop in the accuracy when the image is strongly blurred. The B-Net’s performance is high only for the blur level used in training (σ =4) and blurs of similar strength. B2S-Net did not show much improvement in blur tolerance.

**Figure 3:**
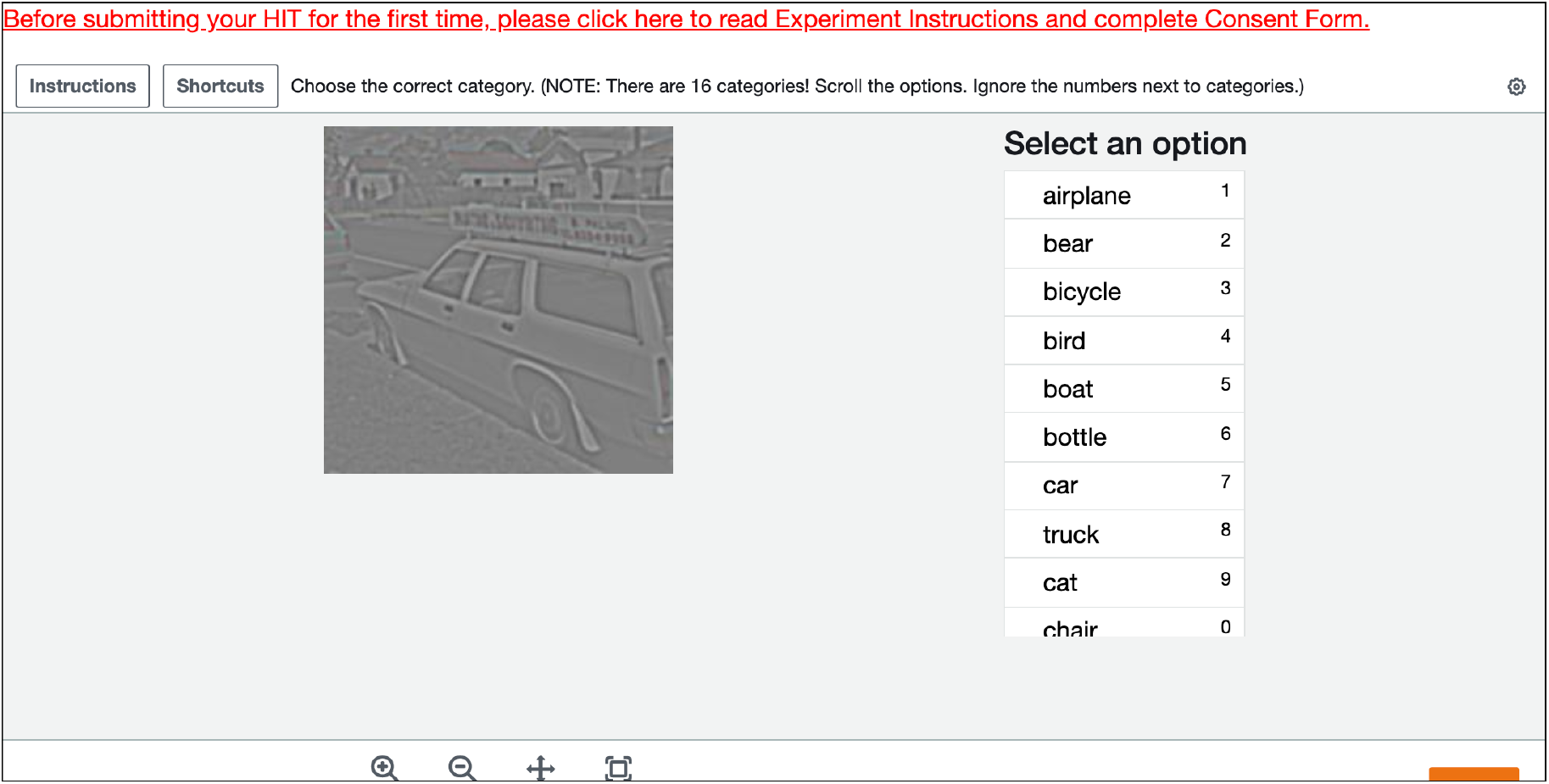
Image classification task on Amazon Mechanical Turk

**Figure 4:**
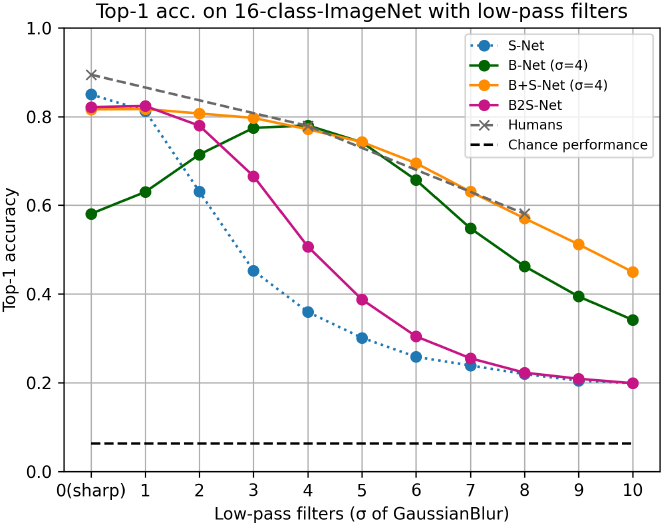
Classification accuracy for low spatial frequency images. The horizontal axis is the standard deviation of the Gaussian kernel (*σ*).

On the other hand, in the low-frequency image recognition test, B+S-Net, which was trained on both blurred and sharp images simultaneously, was able to recognize a wide range of features from sharp images to blurred images of various intensities (blur-robustness). The robustness of the B+S-Net against blur is similar to that of human behavior data.

The B2S-Net showed only a tiny improvement in terms of accuracy over the S-Net, and the B+S-Net showed stronger blur-robustness than the B2S-Net.

### 3.2 Testing robustness to other types of image manipulations

In the previous section, we showed that the models trained on both low-pass-filtered and sharp images acquire robustness to broad range of image blur strengths. Thus, the model might have acquired the ability to capture global features in the image. To gain a more detailed insight into what visual functions were acquired by blur-training, we investigate the behavior of blur-trained models for image manipulations that were not used in the training.

#### 3.2.1 Recognition performance for band-pass images

First, we used band-pass images to investigate the recognition performance of the model in each frequency band. The band-pass images were created by subtracting the two low-pass images of different *σ* (Fig. 5). Using these band-pass images, we were able to find out in which frequency bands Blur-training is influential. We also analyzed whether there is a difference between CNNs and humans regarding the frequency bands they can use for object recognition.

**Figure 5:**
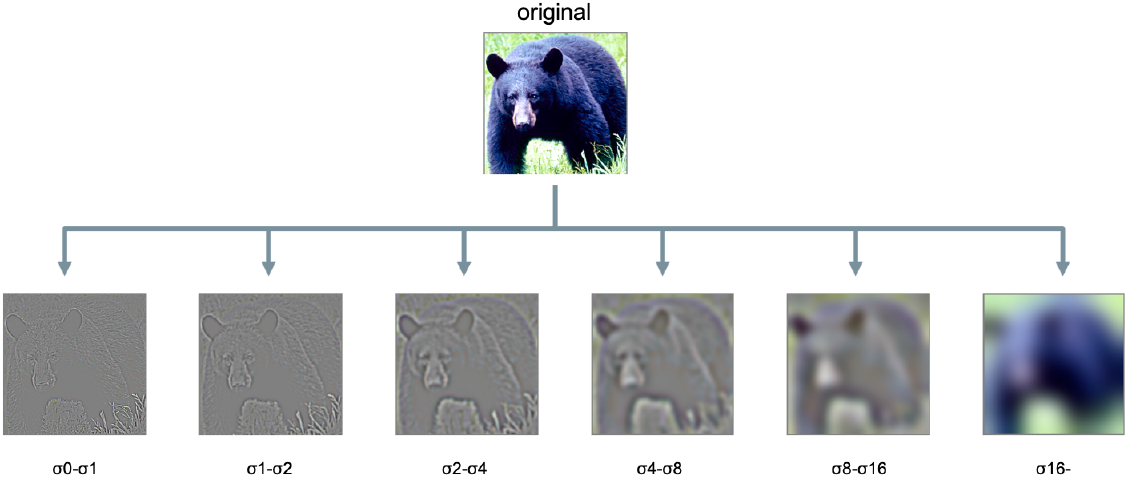
Band-pass images (*σ*(*px*): standard deviation of the Gaussian kernel)

In our experiments, we performed a classification task on band-pass images for CNN models and humans, respectively. In the human experiments, we used AMT to conduct the image classification task for bandpass images *σ*1*px* − *σ*2*px* and *σ*4*px* − *σ*8*px*. The details are described in Section 2.4.

The results (Figure 6) show that B+S-Net improves the accuracy of image recognition over a broader frequency range than the other models, and this indicates that training on blurred images is effective in acquiring the ability to recognize a broader range of frequency features. However, it did not show much effect on images in the high-frequency band.

**Figure 6:**
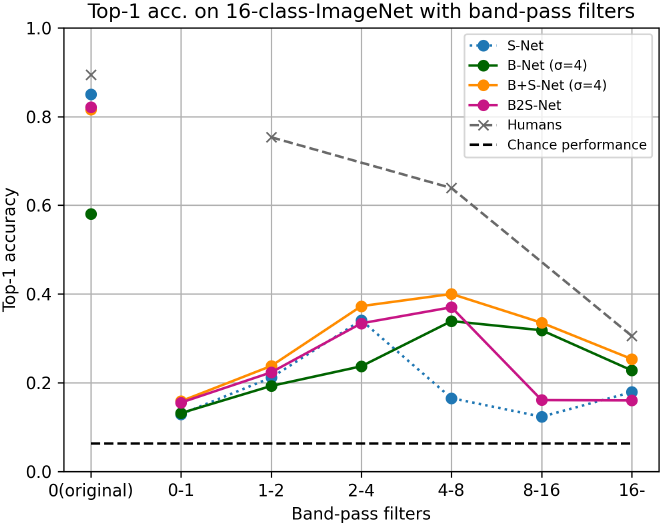
Recognition accuracy for band-pass images. The horizontal axis represents the passband of band-pass images, denoted by *σ* values of the two low-pass-filtered images used for generating the band-pass image.

Next, we compared the accuracy of humans and CNNs. The CNN model showed a lower recognition rate for band-pass stimuli, especially in the high-frequency range.

### 3.3 Recognition performance for global configuration made of local patches

We further investigated whether the blur training could change the global information processing in CNN models by using a test procedure proposed in [Keshvari et al., 2021].

Keshvari et al. [Keshvari et al., 2021] tested the difference in recognition performance between humans and CNNs by manipulating local patches. They divided the original image into several square tiles. There were four partitioning scales: (4 × 4), (8 × 8), (16 × 16), (32 × 32). A Jumbled image was one in which tiles were randomly replaced horizontally, preserving local information but distorting global shape and configural relationships. The Gray Occluder image, in which tiles were alternately greyed out, preserved the global shape and configural information but lost some local information. The Jumbled with Gray Occluder image combined the operations of the Jumbled and Gray Occluder images, and both local and global information were destroyed.

Keshvari et al. [Keshvari et al., 2021] compared the difference in recognition accuracy between a CNN (VGG16 pretrained on ImageNet) and human observers using 640 images from 8 classes in ImageNet (Figure 7 on the left). The CNN showed a significant decrease in accuracy when local information was lost, as in the Gray Occluder image, while it was more tolerant than humans to changes in global configuration when local information was preserved, as in the Jumbled image. On the other hand, humans were more robust than the CNN to the loss of local information as shown in the result of Gray Occluder images, although it was more difficult for them to recognize objects when Jumbling distorted the global shape information. In other words, CNN relies more on local information for object recognition, while humans make good use of global shape information and positional relationships for object recognition.

**Figure 7:**
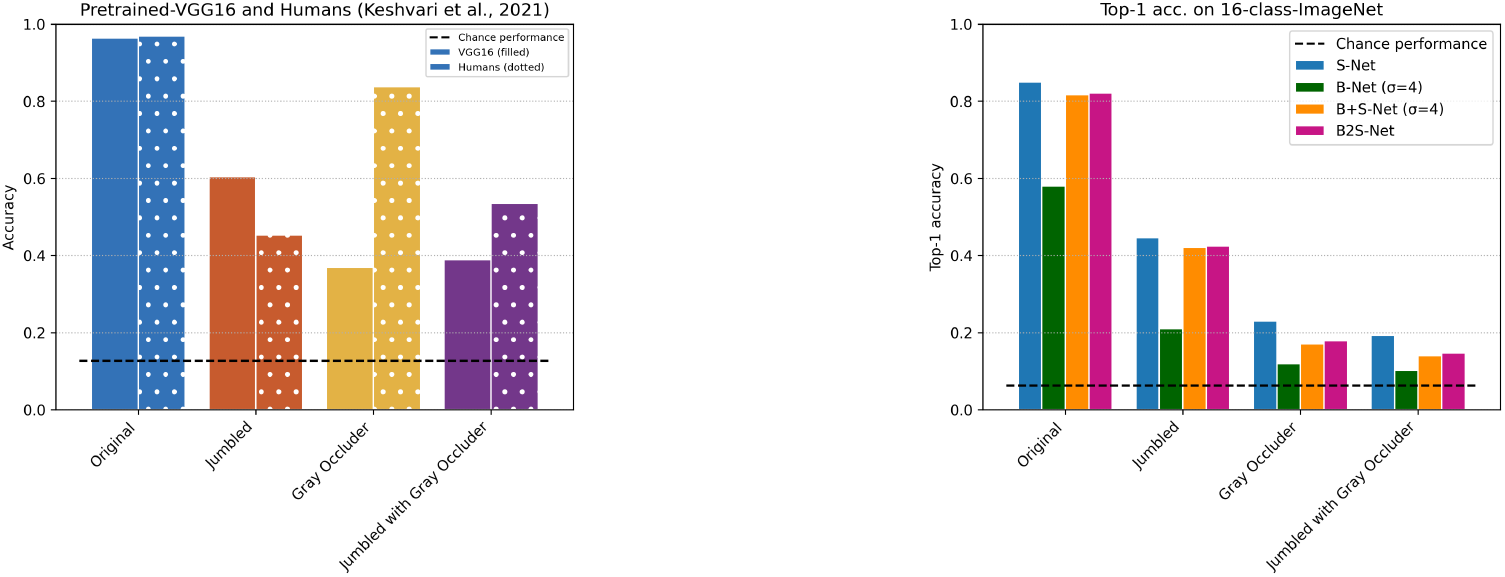
Spatial Shape Recognition. Average percentage of correct answers for (4 × 4), (8 × 8), (16 × 16), (32 × 32). times (Left) Pretrained-VGG16 and human. Data reference [Keshvari et al., 2021]. Test images: 640 images of 8 classes from ImageNet. (Right) 16-class-AlexNet. Test images: 16-class-ImageNet

In this study, we generated the jumbled/occluded images from the test images of the 16-class ImageNet in the same way as in Keshvari et al. [2021] and investigated whether the recognition performance of CNNs becomes closer to that of humans by blur-training (Figure 7 on the right). The results showed that training on blurred images did not change the overall trend of recognition performance on this test set. B+S-Net and B2S-Net did not improve the accuracy for Gray Occluder images compared to S-Net and the accuracy for Jumbled images remained higher than that for Gray Occluder images. The performance drops from Gray Occluder to Jumbled with Gray Occluder were also very small for all the CNN models. These results suggested that the CNN models fail to utilize the global configural information even after the blur-training.

#### 3.3.1 Recognition performance for texture-shape cue conflict images

To investigate whether the blur-trained models show a preference for shape information or texture information, we tested the shape bias proposed by the work of Geirhos et al. [Geirhos et al., 2019].

[Geirhos et al., 2019] created a texture-shape cue conflict image dataset where the texture information of one image was replaced by that of another image in a different class by using the style transfer technique of [Gatys et al., 2016] ^2^. The dataset consists of the same 16 classes as in the 16-class-ImageNet while each image has two correct labels based on its match to the shape or texture class. In total, the dataset contains 1,200 images (75 images per class). The shape bias measures how often the model answers the shape class when it correctly classifies a cue conflict image into either the shape or texture class, and is calculated by the following equation:

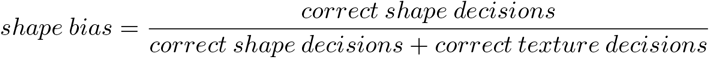

According to the results of Geirhos et al.[Geirhos et al., 2019], while humans showed strong shape bias, CNN models trained on ImageNet showed weak shape bias (in other words, they showed texture bias). When the CNN models were trained on Stylized-ImageNet (SIN) dataset, in which the texture information of an image was made irrelevant to the correct label by replacing the original texture with that of a randomly selected painting, the shape bias of the CNN models (SIN-trained-Net) became closer to human’s level. Moreover, we found SIN-trained-Net has a higher recognition rate for high-pass ans band-pass images as humans do (Figure 8). However, training with SIN is biologically implausible and therefore not helpful in modeling the development of the human visual system.

**Figure 8:**
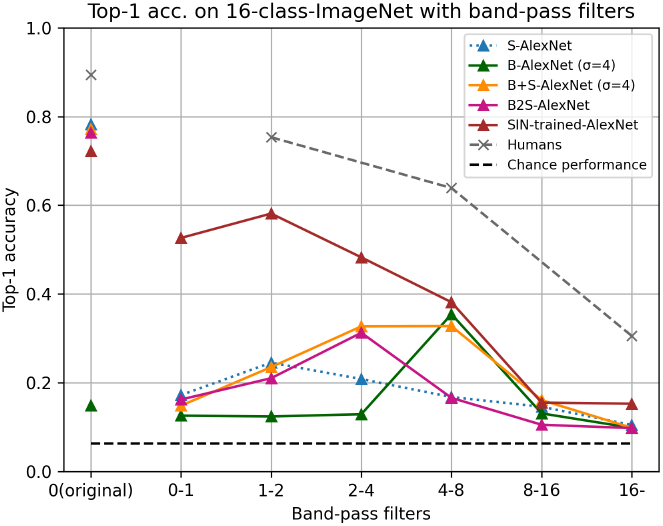
Recognition accuracy for band-pass images. Unlike Figure 6, the Stylized-ImageNet trained network is added, and all CNN models are 1000-class-AlexNet. (The output of the model is determined by extracting 16 classes from 1000 outputs.) SIN-trained-Net [Geirhos et al., 2019]: 1000-class-AlexNet trained on SIN

Here, we calculated the shape bias of the models trained in our study using the texture-shape cue conflict image dataset provided by the authors of Geirhos et al. [2019] to see whether the blur-training could enhance the shape bias of CNNs. Figure 9 presents the shape bias of the four models we trained as well as those of SIN-trained-Net and human data taken from [Geirhos et al., 2019]. Compared to S-Net, B-Net significantly improved the shape bias but decreased the correct classification rate. On the other hand, B+S-Net and B2S-Net improved the shape bias and keep the same classification accuracy as S-Net. However, their shape bias still fell below those of SIN-trained Net and human.

**Figure 9:**
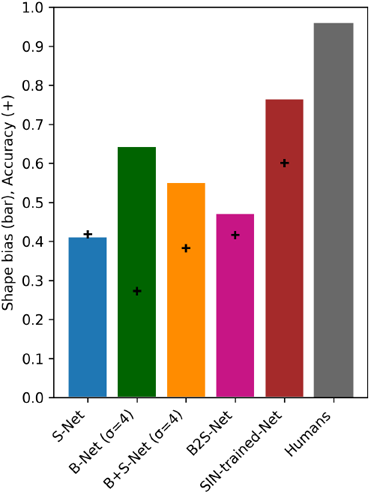
Accuracy (+) and shape bias (bar) for cue-conflict images. SIN-trained-Net[Geirhos et al., 2019]: SIN-trained 1000-class-AlexNet (The output of the model is determined by extracting 16 classes from 1000 outputs.) Humans: data reference [Geirhos et al., 2019]

These results indicate that while training with blurred images has partially succeeded in improving shape bias, it alone is insufficient to bring the bias closer to human level.

### 3.4 B+S-Net with randomly varying *σ*

In the analysis so far, we have fixed the strength of image blur applied to training images for B+S-Net at *σ* = 4. Here, we trained a 16-class B+S-Net while randomly varying *σ* (*σ*0*px* ∼ *σ*4px) and measured its performance on the three test set (Figure 10). As a result, we did not find any significant change in the performance on any of the test set.

**Figure 10:**
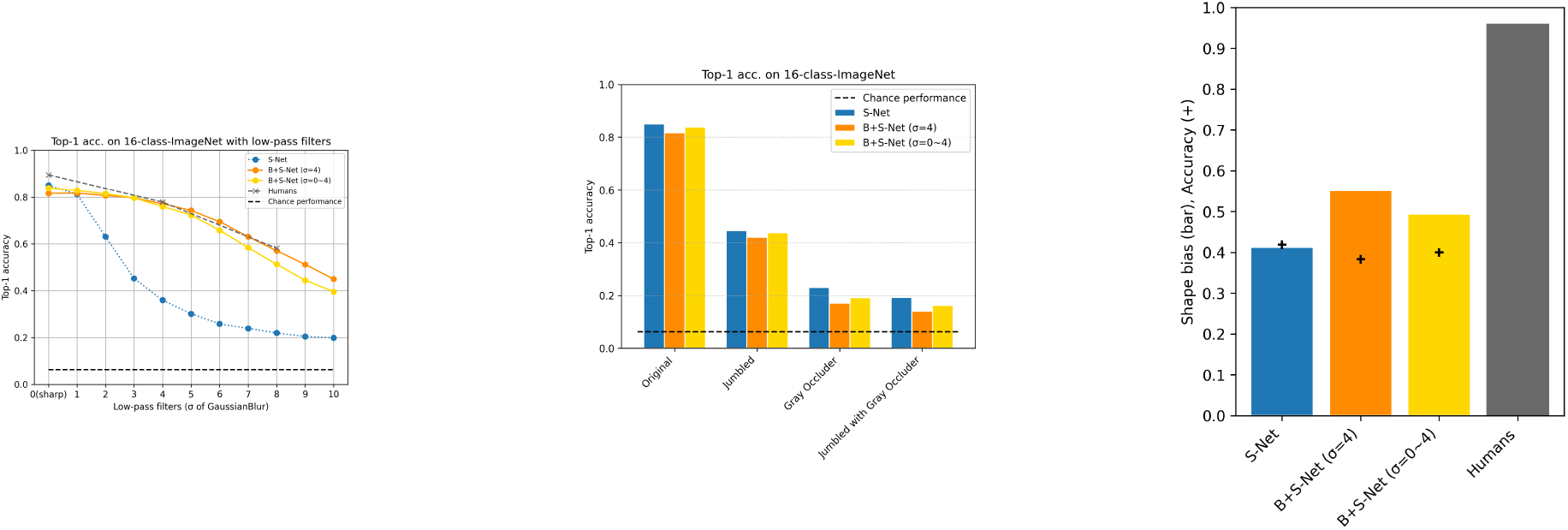
Results for B+S-Net(*σ*0*px* ∼ *σ*4*px*). Left) Low spatial frequency image recognition, Center) Spatial shape recognition, Right) Shape bias

### 3.5 Different model architectures

The analysis so far has been based on the 16-class-AlexNet. We here give an overview of the effects of Blur-training on other network structures.

#### 3.5.1 16-class-AlexNet vs. 1000-class-AlexNet

First, we compare 16-class-AlexNet and 1000-class-AlexNet. The 1000-class-AlexNet has 1000 units in the final layer. The training images are the ILSVRC2012 ImageNet (1000-class classification task, 1.2 million training images), and we trained models from scratch. We trained S-Net, B-Net, B+S-Net, and B2S-Net in the same way as in Section 2.3. As the number of classes and the number of images used in the training increases, it is possible that the 1000-class-AlexNet models learn different features from what 16-class-AlexNet models learn ^3^.

##### Recognition performance for low-pass images

First, we examined the accuracy for blurred images (Figure 11). The 16-class-ImageNet was used to test performances. Therefore, we mapped the output of the 1000-class-AlexNet into 16 classes in the same manner as in Geirhos et al. [2018]. The results of the 1000-class-AlexNet overall exhibited a shared trend with those of the 16-class-AlexNet. B+S-Net showed blur-robustness to a broader range of test blur strengths than S-Net while B-Net showed robustness only around the blur level it was trained with. B2S-Net did not show any improvement over S-Net, probably due to forgetting in the last 20 epochs during which the model was trained with only sharp images. However, we also found that the generalization effect of blur training beyond the blur strength used in the training was smaller for the 1000-class-AlexNet than that for the 16-class-AlexNet. B-Net was firmly tuned to the blur strength used in training (*σ* = 4) and can hardly recognize clear images.

**Figure 11:**
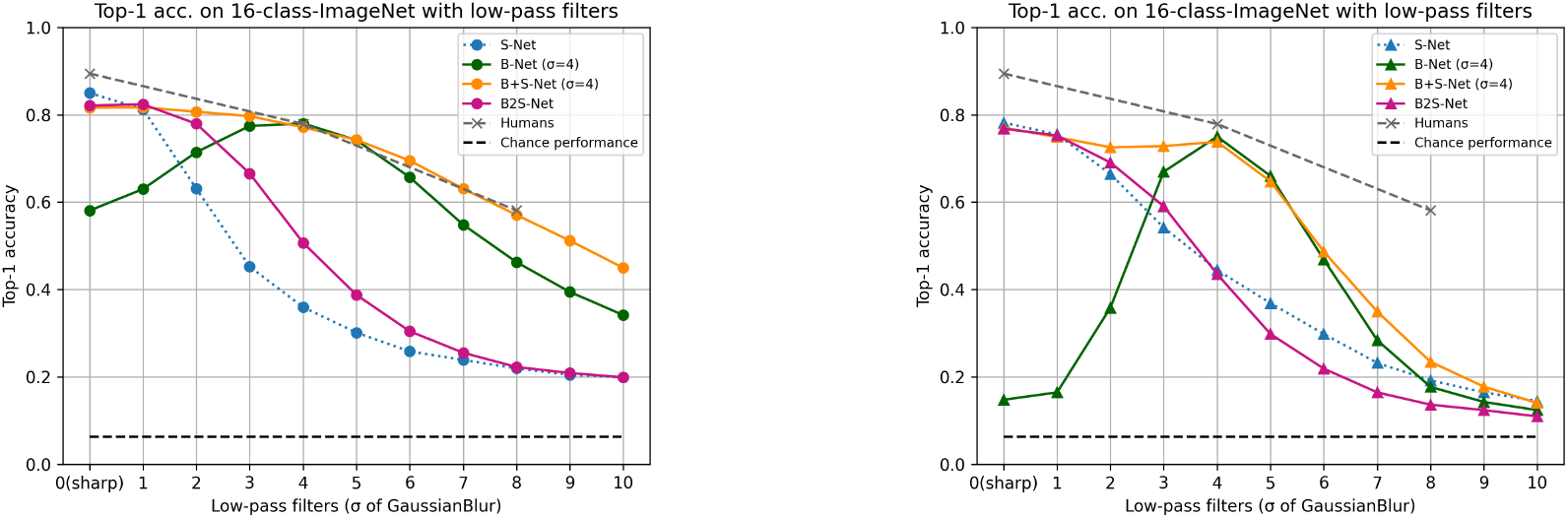
Recognition performance of low spatial frequency features. Left) 16-class-AlexNet Right) 1000-class-AlexNet. (Test image: 16-class-ImageNet)

##### Recognition performance for band-pass images

We also measured the performances on the band-pass-filtered test images (Figure 12). Again, the results of the 1000-class-AlexNet showed a similar trend to the 16-class-AlexNet but the bandwidth that each model could utilize was narrower in the 1000-class version.In addition, as with the 16-class-AlexNet, the accuracy for high-frequency images was low, indicating that the 1000-class models could not recognize the information composed only of high-frequency patterns.

**Figure 12:**
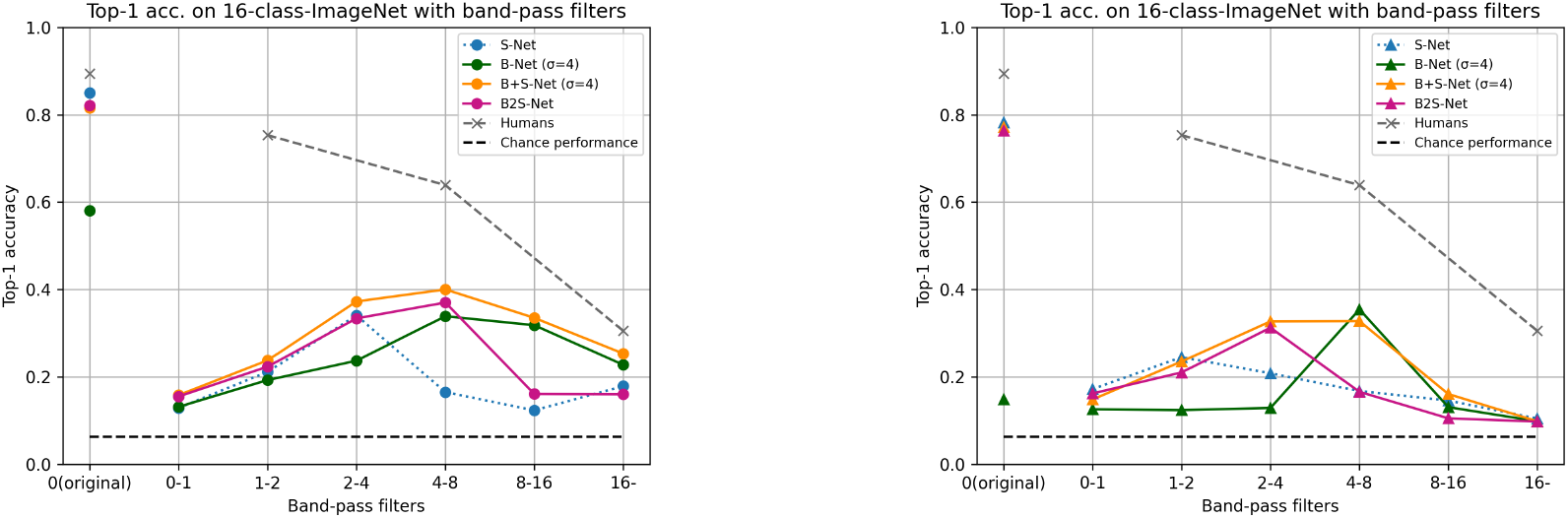
Category classification accuracy for band-pass images. captcha (Left) 16-class-AlexNet. (Right) 1000-class-AlexNet. (Test image: 16-class-ImageNet)

##### Recognition performance for global con figuration made of local patches

We investigated the effect of global configuration of local patches on performances of 1000-class AlexNet. Figure 13 presents comparison of performances between the 1000- and 16-class AlexNet. We found that the trend was exactly the same: high accuracy (about 40%) for Jumbled images and significantly worse accuracy for Gray Occluder images. In other words, the CNNs could classify images to some extent using local information alone, but it was challenging for them to globally integrate the information for object recognition.

**Figure 13:**
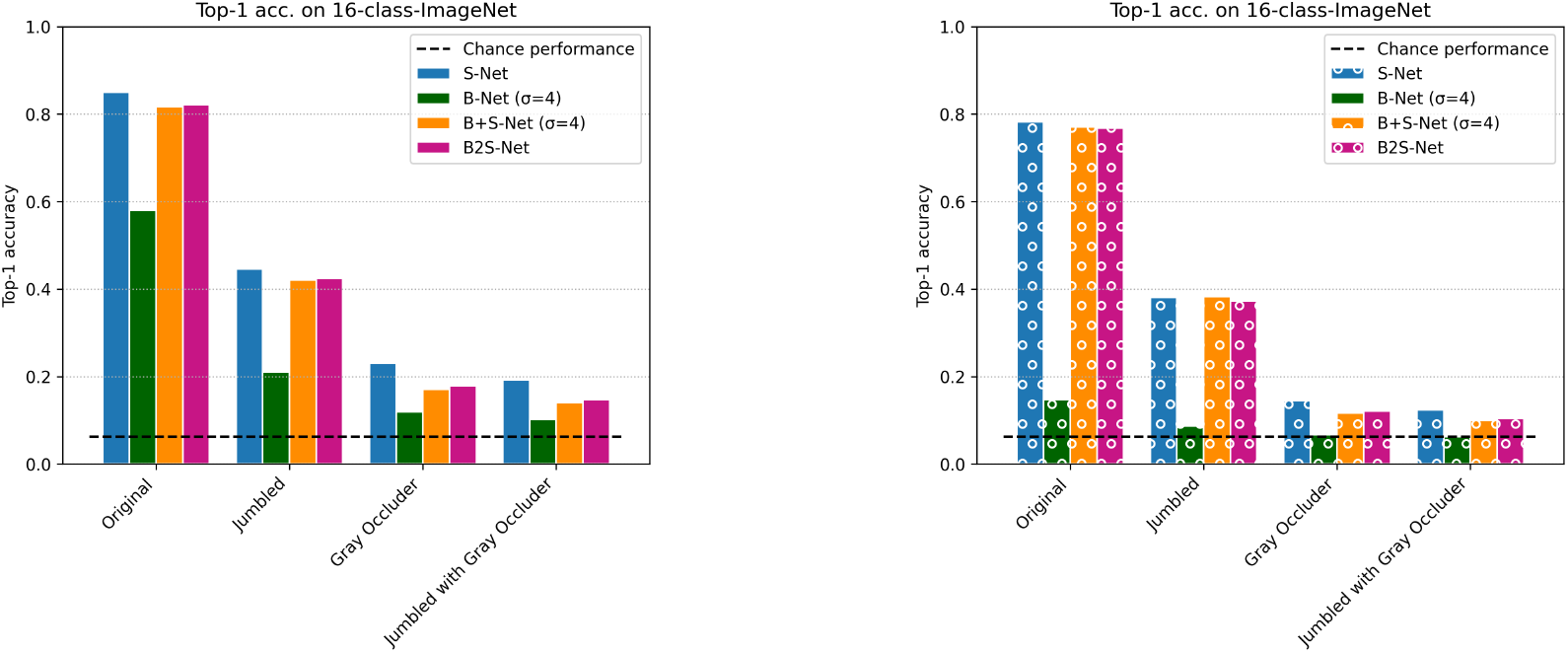
Spatial shape recognition. (Left) 16-class-AlexNet. (Right) 1000-class-AlexNet. (Test image: 16-class-ImageNet)

##### Recognition performance for texture-shape cue conflict images

The results of the shape bias using the cue conflict images (Figure 14) show that there was little effect of blur training on shape bias when the 1000-class dataset was used. However, it should also be noted that the accuracy of the 1000-class models for the cue conflict images themselves was very low, meaning that the models could hardly classify the test images to either the correct shape or texture label in the first place.

**Figure 14:**
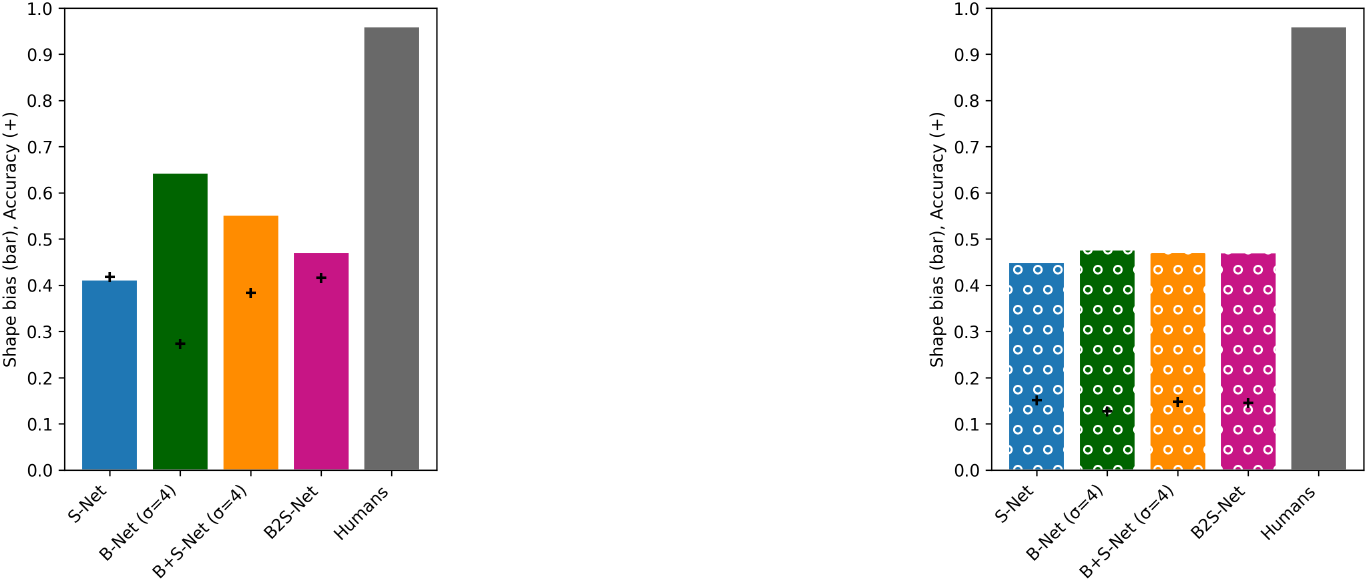
Shape bias. left) 16-class-AlexNet, right) 1000-class-AlexNet

#### 3.5.2 VOneNet (16-class)

In this section, we examine the performance of VOneNet with and without the blur-training.

VOneNet is a model in which the first layer of the 16-class-AlexNet is replaced with a VOneBlock[Dapello et al., 2020]. The VOneBlock is a computational model that simulates the visual information processing in the V1 cortex of the brain, such as the response properties of simple cells and complex cells. It also simulates the stochasticity in neural responses by introducing noise. Importantly, multiscale Gabor filters tuned to low to high spatial frequencies are hard-coded in the VOneBlock. Thus, we expected that the model with VOneBlock might be able to show stronger robustness to low-pass and band-pass filtered images and less reliance on local image features.

Contrary to this expectation, the introduction of the VOneBlock did not change the performance significantly. As shown in Figure 15, the results for each test set presented a remarkable degree of similarity between the models with and without VOneBlock. Thus, changing the lower-level layer to a model closer to the visual cortex did not affect the performance in terms of frequency and shape recognition.

**Figure 15:**
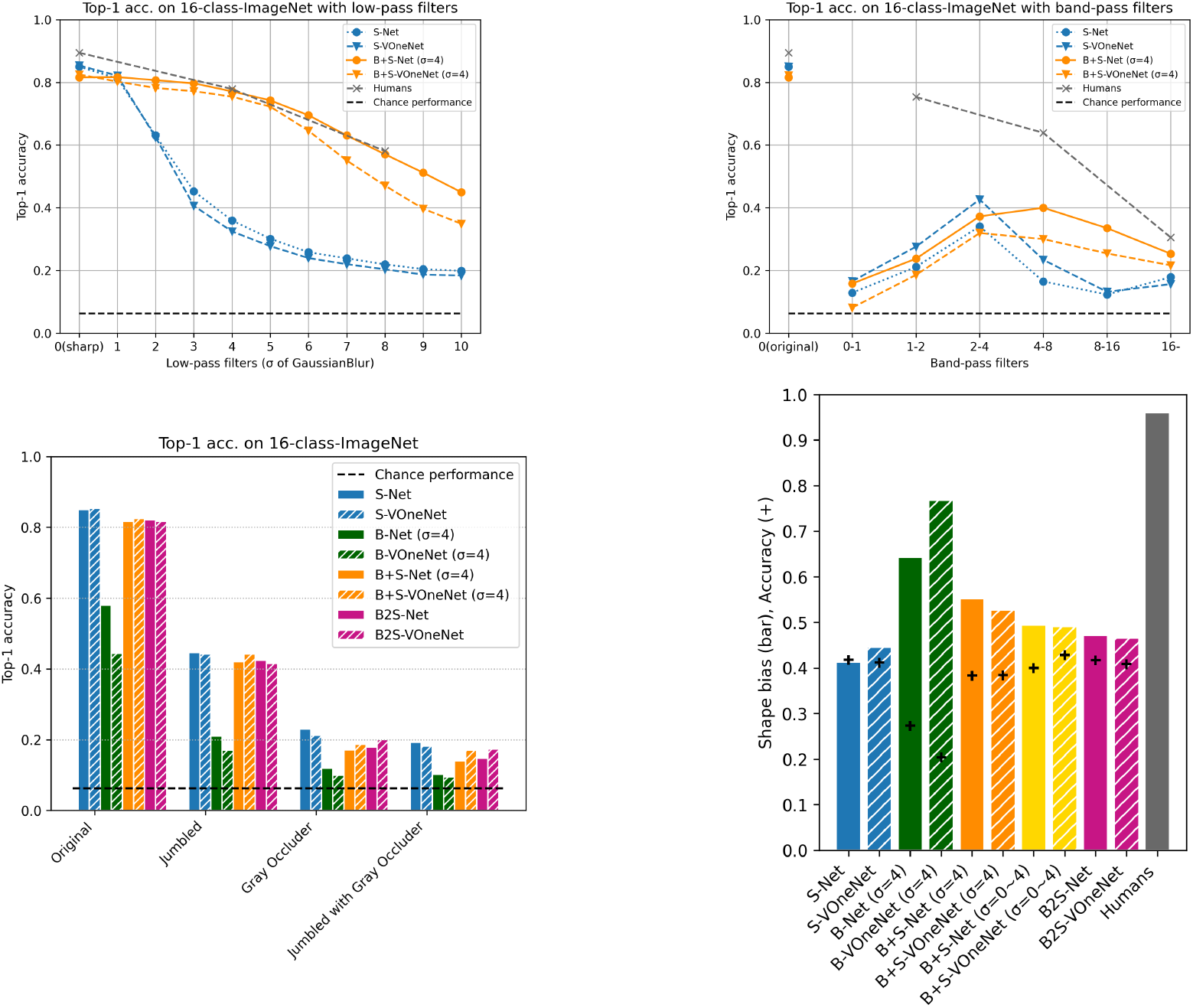
Results for VOneAlexNet. (top left) low spatial frequency image recognition, (top right) bandpass image recognition(bottom left) spatial shape recognition, (bottom right) shape bias

#### 3.5.3 VGG16, ResNet50 (16-class)

In this section, we examine the performance using different network architectures other than AlexNet. The networks studied here are VGG16[Simonyan and Zisserman, 2015] and ResNet50[He et al., 2016]. We set the final output of all networks to 16 classes. We explicitly indicate the name of each network along with the training procedure in this section (e.g. S-AlexNet for S-Net with AlexNet architecture).

##### Recognition performance for low-pass images

The results of performances of the three architectures (AlexNet, VGG16, and ResNet50 from left to right) on low-pass filtered images are presented in Fig. 16). We found the overall trend of results for VGG16 and ResNet50 to be similar to that for AlexNet. However, compared to B-AlexNet, B-VGG16 and B-ResNet50 showed tendency to overfit to the blur strength used during training (i. e., *σ* = 4). Although B+S-VGG16 and B+S-ResNet50 outperformed humans on some low-frequency images, the accuracy dropped sharply around *σ* = 6 and fell below humans at *σ* = 8. In this respect, B+S-AlexNet appears to have the most human-like performance and blur robustness among the three architectures.

**Figure 16:**
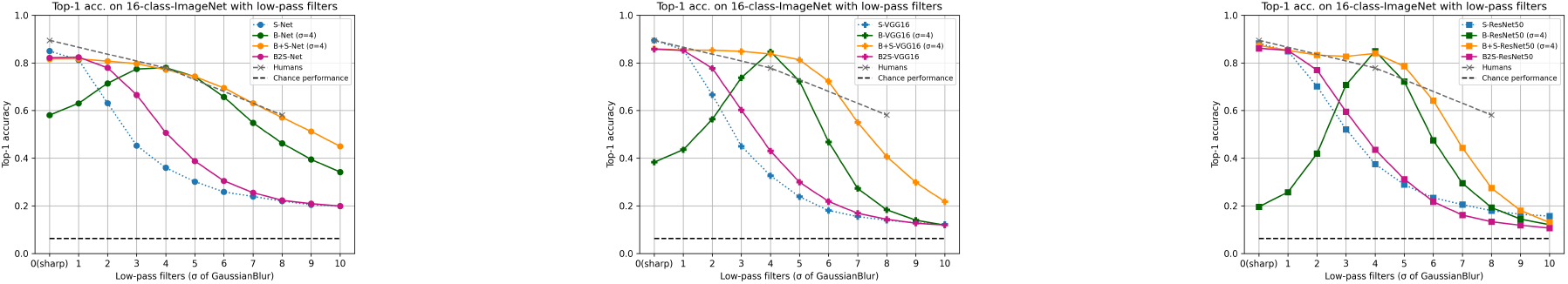
Correct classification rates for low spatial frequency images: left) 16-class-AlexNet, middle) 16-class-VGG16, right) 16-class-ResNet50

##### Recognition performance for band-pass images

Next, we compare the accuracy for the band-pass images. The results of the three architectures are presented in Fig. 17. Although variation in tuning patterns due to training methods was similar across architectures, we found that VGG16 (in particular, S-VGG16) was better at recognizing objects using high frequency components than the other architectures. Training with blurred images decreased the reliance on high frequency components in VGG16.

**Figure 17:**
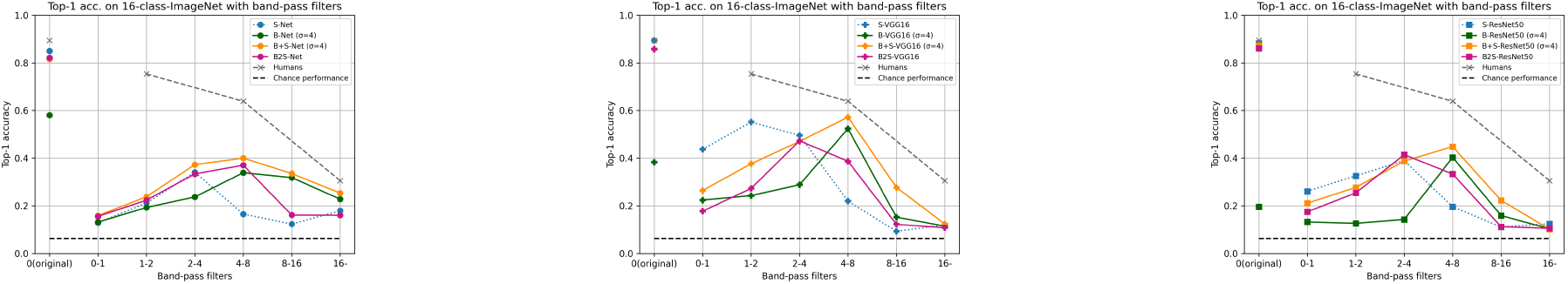
Classification accuracy for bandpass images. left) 16-class-AlexNet, middle) 16-class-VGG16, right) 16-class-ResNet50

##### Recognition performance for global configuration made of local patches

The results of the local patch jum-bling/occlusion test showed no significant difference among the training methods in all three architectures (Figure 18). Regardless of the architecture, the CNN models tended to rely on local information rather than global configural information for object recognition.

**Figure 18:**
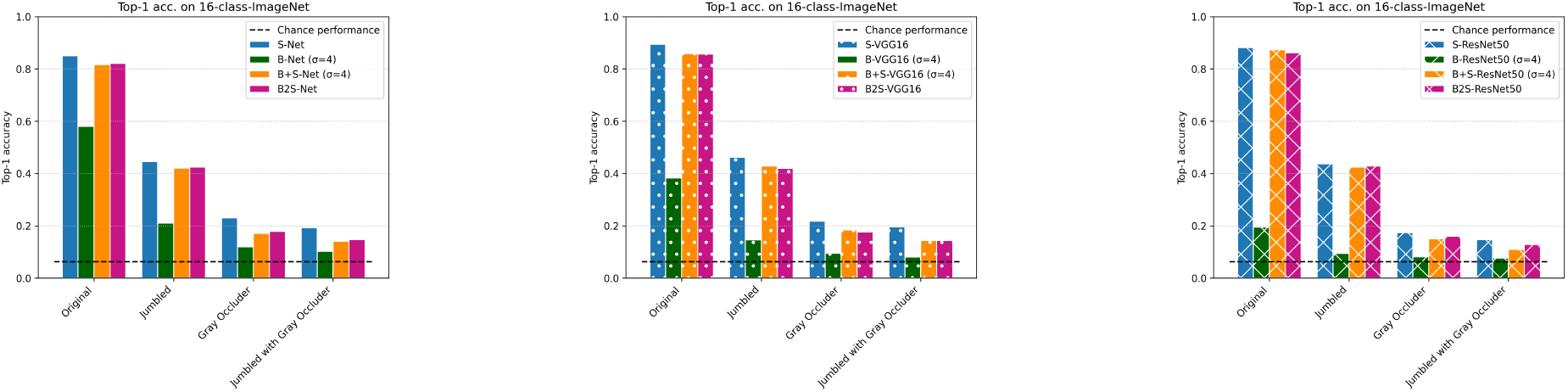
Global/Local recognition performance. left) 16-class-AlexNet, middle) 16-class-VGG16, right) 16-class-ResNet50

##### Recognition performance for texture-shape cue conflict images

The shape bias of the three architectures are presented in Fig. 19. In all architectures, the training with blurred images increased the shape bias. It should be also noted that the shape bias was lower (the texture bias was higher) for S-VGG16 and S-ResNet50 than for S-AlexNet. This result may be related to the fact that S-VGG16 and S-ResNet50 had higher accuracy for high-frequency images than S-AlexNet, as shown in Figure 17. Thus, decreasing the reliance on high frequency components might have some effect in increasing the shape bias. However, the increase in shape bias due to blur-training was not enough to reach the human level in any of the three architectures.

**Figure 19:**
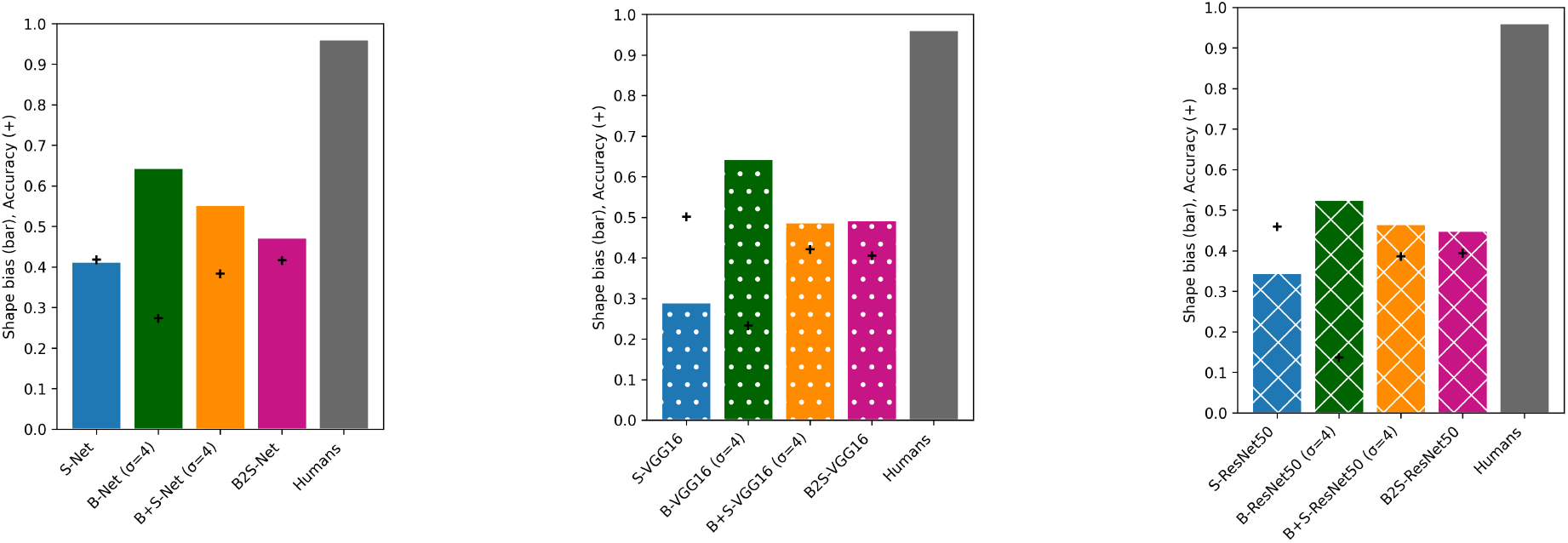
Classification accuracy for band-pass images. left) 16-class-AlexNet, middle) 16-class-VGG16, right) 16-class-ResNet50

### 3.6 Summary and Discussion of Section 3

In Section 3, we compared the effect of training with blurred images on the CNN models in terms object recognition performance. In all CNN models we tested, the recognition performance of low spatial frequency features improved by the blur training. In particular, the model trained simultaneously on blurred and sharp images (i.e., B+S-Net) showed blur robustness against a wide change in image blur close to that of humans. We will further analyze the mechanism behind this similarity in the next section (Section 4).

The blur-robustness of B2S-Net was weaker than that of B+S-Net. In other words, the models showed better performance when trained on blurred and sharp images simultaneously, rather than on a schedule that simulated the human visual development. This result apparently disagrees with the study of Vogelsang et al.[Vogelsang et al., 2018], which showed that training in the order of low resolution to high resolution improved blur robustness of a CNN model in face recognition. This difference can be attributed to the difference in the task adopted in our study and Vogelsang et al.’s (i.e., general object classification v.s. face classification) [Jang and Tong, 2021]. A recent study using object recognition [Avberšek et al., 2021] reports the effect of training schedule consistent with ours. The task difference may be related to the fact that the optimal discriminative features for object recognition are biased toward high frequencies while only low-frequency features are sufficient for good face classification accuracy [Jang and Tong, 2021].

The failure of B2S-Net to recognize blurry images indicates that simply simulating the development of visual acuity during training cannot account for the blur robustness of human vision in general object recognition. However, even after the completion of visual development, we still experience blurred retinal images on a daily basis due to defocus as well as motion blur, and scattering caused by climatic conditions such as rain and fog and by the transmission of translucent objects. In this sense, B+S Net, trained simultaneously with both blurry and sharp images, can be regarded as reproducing biologically plausible situations to some extent. In addition, the CNNs we tested do not include a mechanism that prevents the forgetting of previously learned representations. The B2S-Net may have forgotten the processing for low-frequency components because it was trained only on sharp images in the last 20 epochs. Therefore, B2S-Net may be able to recognize blurred images as well as B+S-Net by adding a mechanism that prevents the model from forgetting the representations tuned for blurred images learned in the early phase of training. Machine learning literature has suggested a few methods to prevent so-called catastrophic forgetting in continual learning [de Melo et al., 2022]. One is to protect the weights relevant to the stimuli learned in the early phases of training [Kirkpatrick et al., 2017]. This reminds us of the critical period of the biological neural networks. Another method is to use the memory of relevant prior information to retrain the network with new information [Aljundi et al., 2019]. This mechanism will make the effect of B2S training quite similar to that of B+S training.

Our results also show that the B+S Net that has acquired human-level blur robustness has not acquired human-like global visual processing. The performance test using band-pass filtered images showed that all CNN models, including B+S Net, were not good at utilizing band-limited features while humans retained good accuracy in the mid-to-high frequency range. Although the shape bias of the blur-trained models was slightly enhanced, it was not enough to reach the human level. The test using the jumbled patch images and images with local occlusions revealed that all the models relied mostly on local features, did not utilize the global configuration, and were critically vulnerable to local occlusions. All these results are in stark contrast to human visual processing, which is known to rely more on global configural relationships and shape information and is less sensitive to partial occlusions in object recognition tasks. Therefore, the results obtained in this section indicate that the information processing learned in B+S-Net is still quite different from that of the human visual system.

Our results reveal that what the networks cannot acquire from blur training is human-recognizable global configuration features present in high-pass images and texture-shape cue conflict images. Note that the similarity of these classes of images is supported by a finding that the SIN-trained Net shows good recognition for high-pass images as well. In high-pass images, local edge features defined by high-frequency luminance modulations produce global configurations at a scale much larger than a fine-scale edge detector. For detection of these global features, second-order processing such as those modeled by an FRF (filter-rectify-filter) model for human vision (e.g., Graham and Landy [2002]) may be necessary. It seems that object recognition training with sharp and blurred images alone does not provide neural networks with the ability to process second-order features.

According to the comparison of the model architectures, there was no qualitative difference in the effect of Blur-training. Importantly, we found that VOneNet, that hard-coded the computational processes in the primary visual area (V1) in the front-end of AlexNet, did show no improvement in any tasks tested in this study. This indicates the limited impact of the initial layer on the frequency tuning at the task performance level and on the mid-to-high level information processing related to the shape bias and the configural effect. On the other hand, we also found a few notable differences in the frequency tuning patterns between the architectures. For example, the loss of blur robustness observed in B2S-Net was more prominent in 1000-class AlexNet as well as in 16-class VGG16 and 16-class ResNet50 than in 16-class AlexNet. B+S-Net and B-Net in these architectures were also more narrowly tuned to the blur strength used during the training. For the 1000-class AlexNet, the reason for this may be attributed to the fact that the models were exposed to a larger number of images (and thus went through a larger number of weight updates) when using the 1000-class dataset than the 16-class dataset. For the 16-class VGG16 and 16-class ResNet50, differences in model architecture such as increased depth, reduced kernel size, and residual connections (in the case of ResNet) may have resulted in improved learning efficiency, thereby making them more likely to specialize in features that are optimal for the current blur strength. In addition, we also found that VGG16 shows a higher accuracy for the band-pass filtered images with high spatial frequency than the other architectures, though we have not yet figured out why.

## 4 Analysis of the internal representation of B+S Net

So far, we have analyzed the effect of training with blurred images on the basis of recognition performance. As a result, we found that the recognition performance of B+S-Net for low spatial frequency images is similar to that of humans. In this section, we try to understand whether B+S-Net processes information similarly to humans through internal representation analysis.

How the information is processed inside the CNN for the sharp and blurry features can be roughly categorized into two patterns.

**Case 1** Representations for sharp and blurry images are integrated in the early to middle stage of the visual processing, after which shared information processing is used (Fig. 20, top).

**Figure 20:**
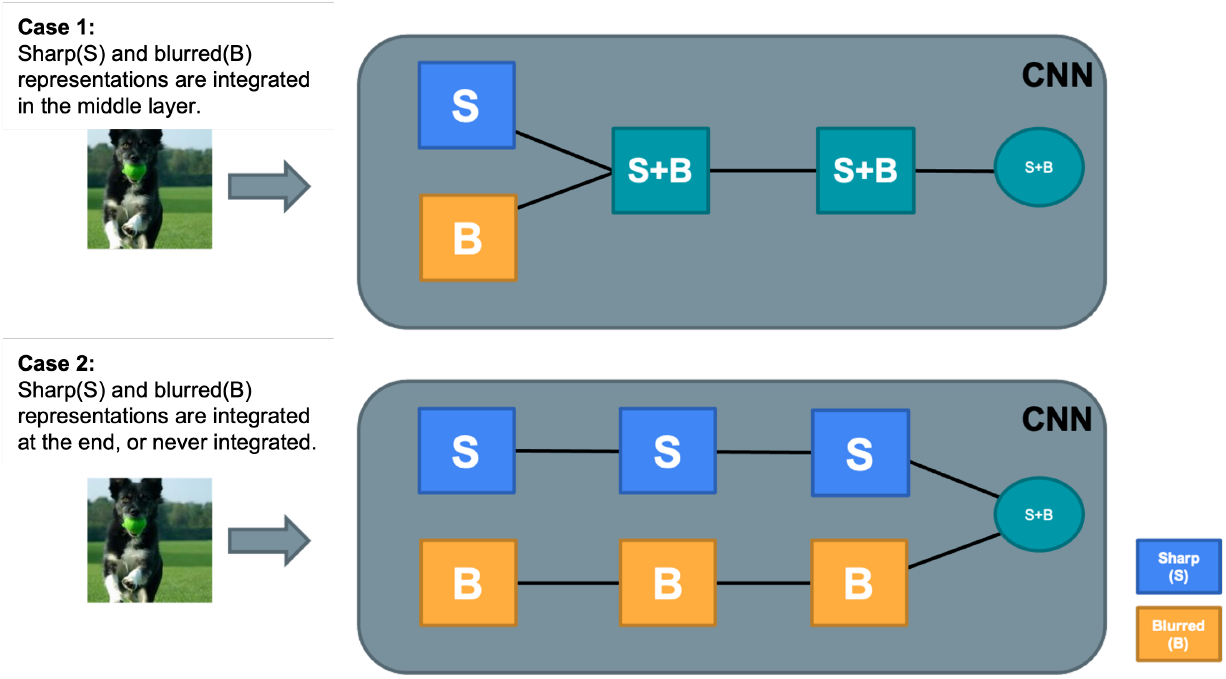
Hypothesis of intermediate feature representation and information processing

**Case 2** The sharp and blurred image features are processed separately and integrated in the last stage to recognize the object. (Fig. 20, bottom).

We expect that robust recognition systems, probably including humans, should process sharp and blurred images as in case 1. On the other hand, CNNs with a powerful learning ability may create specialized sub-network, each processing blurred images and sharp images separately, to optimize the performance. To use CNNs as a computational tool to understand human-like robust processing, we should check whether the processing strategy to achieve blur-robustness is not very different between humans and CNNs.

### 4.1 Frequency tuning of filters in the first layer

First, we visualized the receptive field of the first convolutional layer (Fig. 21), as previous studies did [Vogelsang et al., 2018, Jang and Tong, 2021]. B+S-Net shows a shift of the spatial frequency tuning to the lower frequency side compared to S-Net. In other words, B+S-Net extracts more low-frequency information in the first layer. How this affects the internal representation of low spatial frequencies is discussed in the next section.

**Figure 21:**
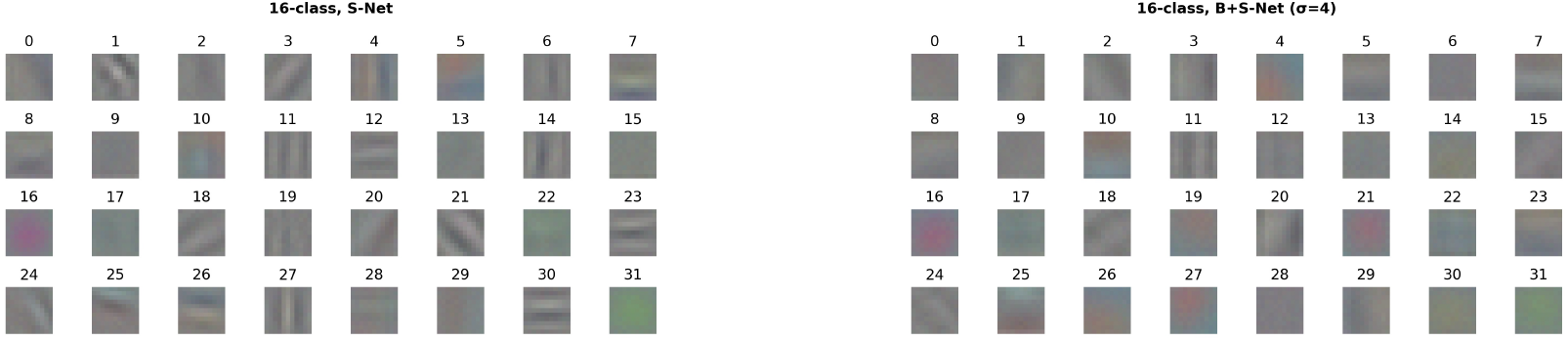
16-class-AlexNet visualization of the first convolutional layer receptive field

Additionally, in the case of 1000-class-AlexNet, features with higher spatial frequencies are extracted (Fig. 22). This may be because they needed to extract finer local features to perform more fine-grained classification.

**Figure 22:**
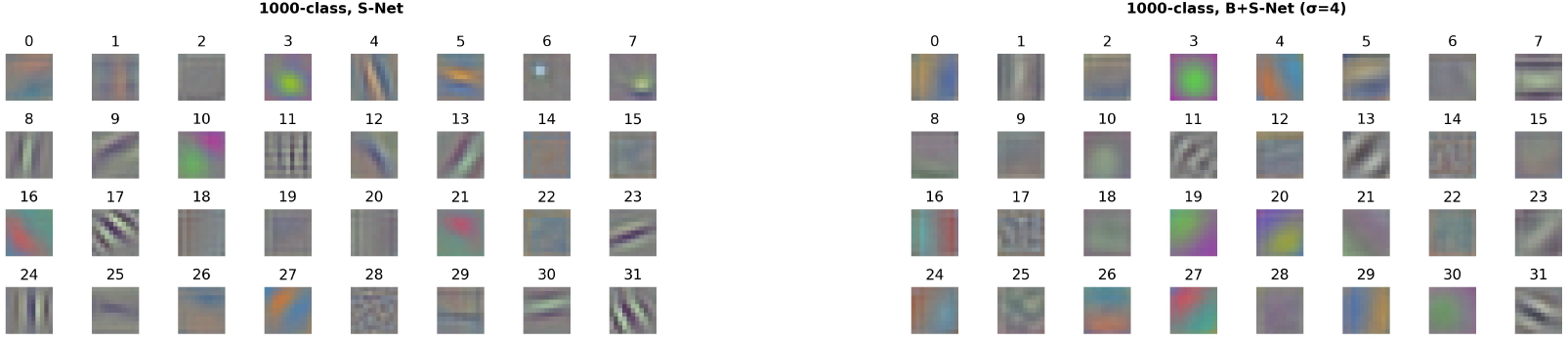
1000-class-AlexNet visualization of the first convolutional layer receptive field It shows each of the 32 out of 64 first convolutional layer filters for each model.

### 4.2 Are sharp image features and blurred image features either combined or separately processed?

#### 4.2.1 Correlation of activity in the intermediate layers

Next, we analyze how the sharp and blurred image features are processed in the intermediate layers of the CNN models. For this, we computed the average correlations of unit activities in each intermediate layer between the sharp and blurred image inputs (S-B correlation). If the processing is shared between the sharp and blurry images, the correlation of activities in the intermediate layer should be high. Here, we calculated the S-B correlations in the following three cases: the sharp and blurry image pair is generated from (1) the same image, (2) different images from the same class, and (3) different images from different classes. For each case, we computed correlations for all possible sharp-blurry image pairs from 1,600 test images of the 16-class ImageNet. Then, the correlations were averaged across image pairs. The unit activities after the ReLU activation function were used to compute the correlations.

Figure 23 presents the S-B correlation in each layer of the 16-class S-AlexNet (left) and B+S-AlexNet (right). When the sharp-blurry image pairs from the identical images are used for input (the blue solid line), S-B correlations are high in the initial layer for both S-Net and B+S-Net. In S-Net, the correlation drops once in the intermediate convolutional layers, and rises slightly again in the fully-connected (FC) layers. On the other hand, in B+S-Net, the correlation remains consistently high from the initial layer to the output layer. This result suggests that the representations for sharp and blurry images are shared from the very early stage in the B+S-Net while they become progressively divergent in the intermediate convolutional layers in the S-Net. When the sharp-blurry image pairs from different images in the same class are used for input (the orange broken line), the S-B correlation remains low through all the convolutional layers for both S-Net and B+S-Net. After that, in both networks, the S-B correlation gradually increases through the FC layers, but in the last layer, it falls at a lower level for S-Net than for B+S-Net. When the sharp-blurry image pairs from different images in the different class are used for input (the green dotted line), the S-B correlation remains low throughout all layers for both S-Net and B+S-Net. The low correlations between convolutional layer activities from different images of the same class (the orange broken line) and the different class (the green dotted line) conditions are because image features are represented in spatially localized manner in the convolutional layers. The increasing correlation in the FC layers in the same class condition indicates that these local features are integrated to obtain position-invariant representations convenient for classification task. The lower correlation in the output layer of the S-Net than that of the B+S-Net matches the results of the performance comparison observed in the previous section.

**Figure 23:**
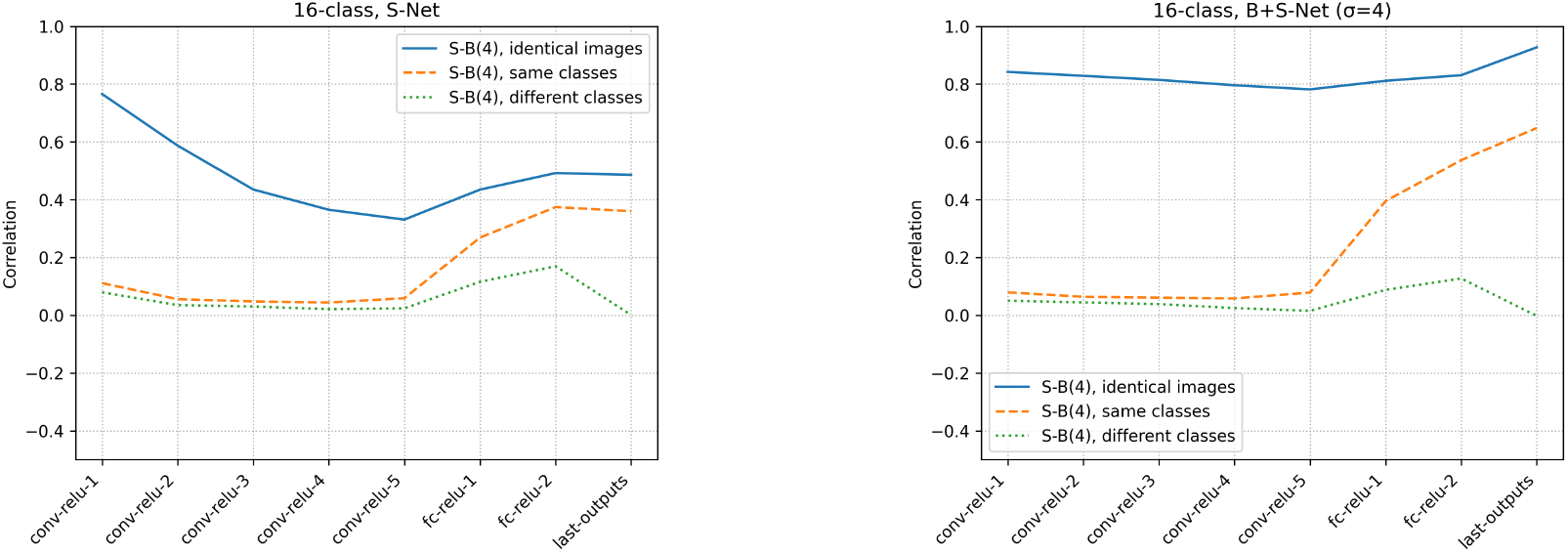
16-class-AlexNet mean correlation of activity in the middle layers units for sharp (S) and blurred (B) images identical images: mean activity correlations for identical images, same classes: mean activity correlations for images of the same class other than identical images, different classes: mean activity correlations for images of different classes

We also applied the same analysis for 16-class VOneNet and 1000-class AlexNet and confirmed that the B-S correlations for both of the network architectures are similar to those we found for 16-class AlexNet, as shown in Fig. 24 (16-class VOneNet) and Fig. 25 (1000-class AlexNet).

**Figure 24:**
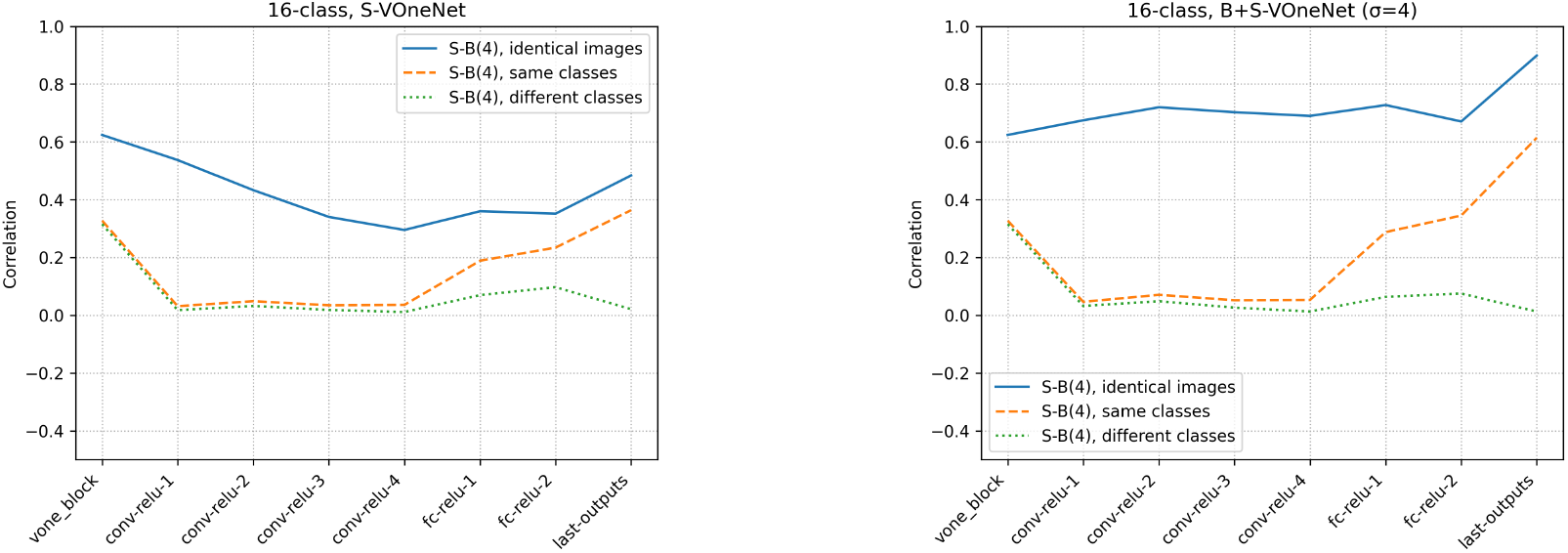
16-class-VOneNet mean activity correlation of the middle layer units for the sharp (S) and blurred (B) images

**Figure 25:**
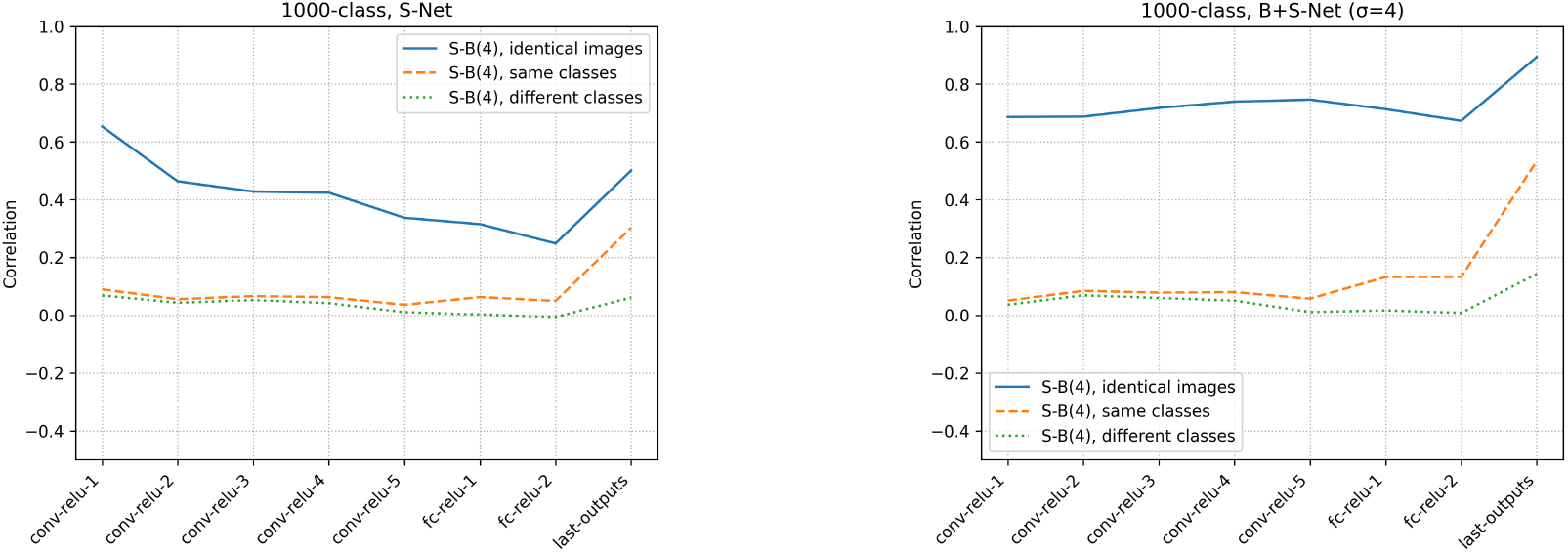
1000-class-AlexNet mean activity correlation of the middle layer units for sharp (S) and blurred (B) images

In B+S-Net, the B-S correlation of the identical image pairs in the intermediate layers remains consistently high at around 0.8. The reason for this could simply be that the high frequency components are cut off by filters in the initial layer, and only low frequency features are processed from the beginning. Indeed, the filters of the first layer show a tendency towards low-frequency preferences (Figure 21). However, the influence of filters in the initial layer is considered to be minor because the difference in the S-B correlation in the initial layer between the S-Net and B+S-Net is small. This is also confirmed by the S-B correlation in VOneNet (Figure 24), in which the first layer is hard-coded as a Gabor filter bank and thus has the same correlation values between the S-Net and B+S-Net. In VOneNet, the difference between the S-Net and B+S-Net widens after the second layer, with the correlation increasing in the B+S-Net and decreasing in the S-Net. This indicates that the B+S-Net forms the features common to both sharp and blurred images from the multiband information extracted in the initial layer.

#### 4.2.2 Visualization of the internal representation by t-SNE

We attempted to understand how the sharp and blurry images are represented in the intermediate layers of the CNN models by visualizing them using the dimensionality reduction algorithm, t-SNE[Laurens van der Maaten and Geoffrey, 2014]. Specifically, we recorded the activities of each layer obtained from sharp and blurry images and compressed them into two dimensions for visualization. The two input parameters of t-SNE, perplexity and iteration, were set to 30 and 1000, respectively. The results shown here are visualizations of 10 pairs of sharp and blurred images of the same image sampled for each of the 16 classes.

The visualizations of the intermediate layer activities for the sharp and blurred images (Figure 26) show that the representations of the sharp and blurry images in the convolutional layers overlap in both S-Net and B+S-Net models, making it difficult to see the difference. However, the representations of the sharp and blurry images in the FC layers become closer in B+S-Net than in S-Net.

**Figure 26:**
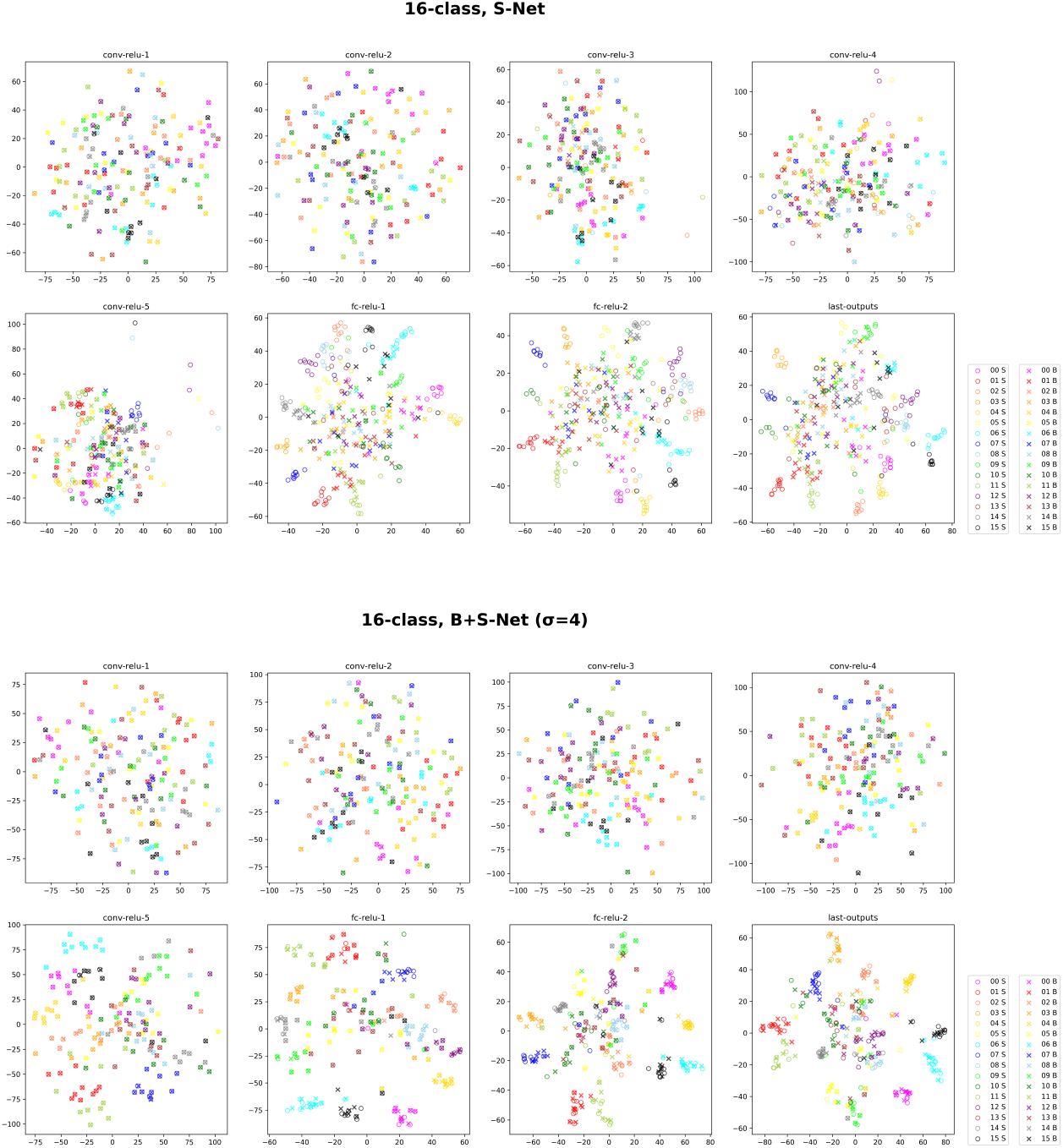
Visualization of the internal representation with t-SNE (sharp image (S) and blurred image (B))

#### 4.2.3 Zero-shot transfer learning

The results of the S-B correlation analysis in Sections 4.2.1 and 4.2.2 suggested that the representations are shared between the sharp and blurry images in the intermediate layers of B+S-Net.

Here, we test the generalizability of the shared representation acquired by the blur training to the unseen classes. In other words, we are interested in whether the blur robustness extends its effect beyond the classes used in the blur training.

For this purpose, we trained a subset of object classes, either one or eight in 16 classes, without using blurry images while training the remaining classes using both blurry and sharp images, and later evaluated the classification accuracy for that subset of classes using blurry images. Conversely, we also trained a subset of classes without using sharp images while training the other classes using both blurry and sharp images, and later evaluated the classification accuracy using sharp images.Therefore, there were in total four conditions, i.e., training without blurry or sharp images for one or eight classes (w/o 1/16B, w/o 8/16B, w/o 1/16S, w/o 8/16S).

The recognition accuracy for the unseen image types (either blurry or sharp) in the four test conditions are shown in Table 2. The confusion matrices of the individual models are also shown in Figs 27, 28, 29 and 30. When either blurry or sharp images were excluded for half of the training classes (Figs. 28 and 30), the models were not able to recognize these classes of images with the unseen image type. On the other hand, when either blurry or sharp images were excluded for one training class (Figs. 27 and 29), the accuracy for the unseen class is not very high, but much higher than the chance level 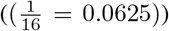. Therefore, although the effect of the generalization of the sharp and blurry features to unseen categories was limited in terms of the zero-shot transfer performance, the small amount of transfer were clearly observed when there was only one excluded class.

**Table 2:** Classification accuracy for B+S-Net trained without using blurred/sharp images for specific object classes. (Average accuracy for multiple labels. Chance 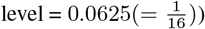

| Training method | Accuracy for untrained labels |
| --- | --- |
| Training without B images for one class (w/o 1/16B) | 0.19 |
| Training without B images for half of the classes (w/o 8/16B) | 0.05 |
| Training without S images for one class (w/o 1/16S) | 0.15 |
| Training without S images for half of the classes (w/o 8/16S) | 0.03 |

**Figure 27:**
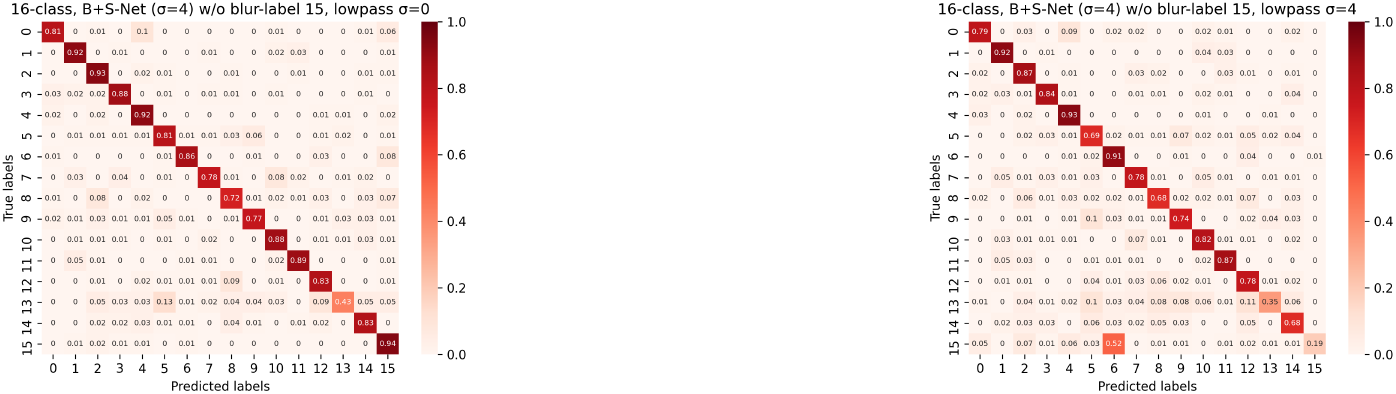
Confusion matrix when one blurred label (No. 15) is excluded from the training

**Figure 28:**
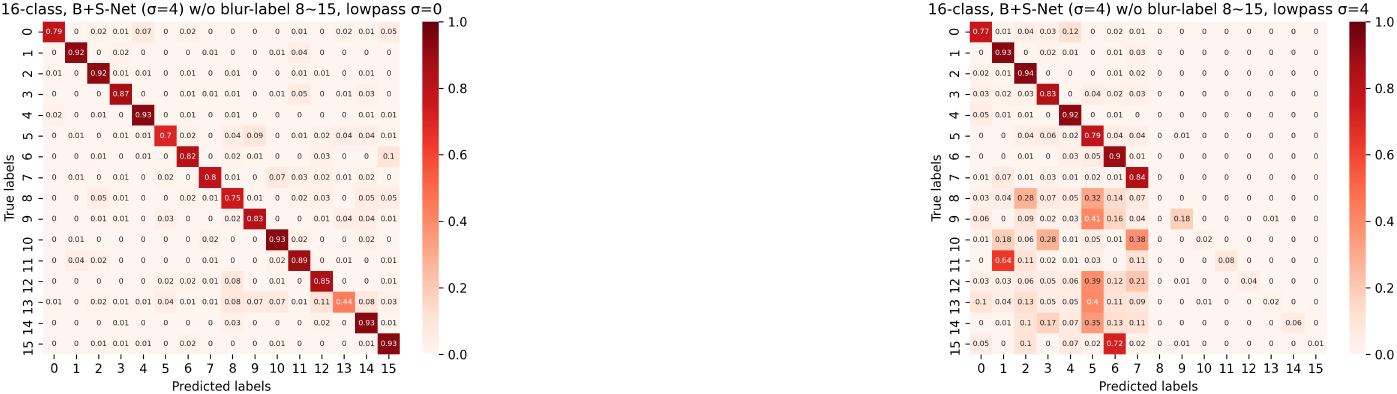
Confusion matrix when half of the blurred labels (No. 8-15) are excluded from the training

**Figure 29:**
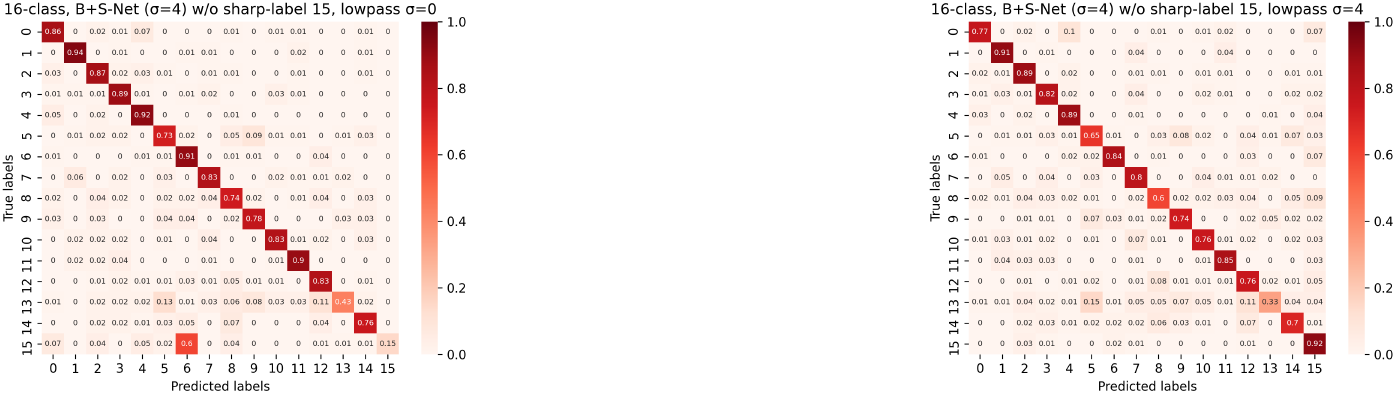
Confusion matrix when one sharp label (No. 15) is excluded from the training

**Figure 30:**
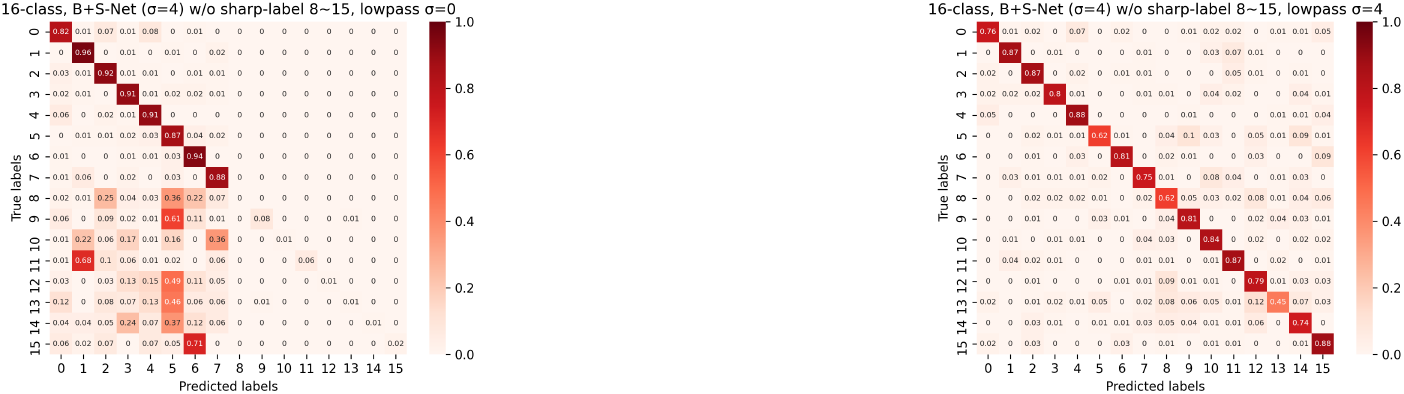
Confusion matrix when half of the sharp labels (No. 8-15) are excluded from the training

To further analyze the internal representations of the models trained in the transfer experiment, we examined the S-B correlations in the intermediate layers of each model. As a result, we found that in the model that did not use one of the image types for training the eight classes (the right-hand side of Figs. 31 and 32), the S-B correlation from the identical images is significantly reduced in the middle to high layers for the unseen category (blue line), compared to that for the seen category (orange line). On the other hand, in the model that did not use one of the image types for training the only one class (the left-hand side of Figs. 31 and 32), the S-B correlation from the identical images for the unseen category (blue line) remains almost as high, albeit slightly lower than that for the seen category (orange line). Therefore, although the shared representation for the blurry and sharp images did not seem to generalize well to the unseen class in the performance level, the similarity of the internal representations appeared to be high between the seen and unseen classes for the model with one excluded class. The reason for this apparent discrepancy is presumably because the misclassification to a class with a similar representation can still occur due to the imperfect alignment of blur-sharp representations. In fact, the misclassifications in the model with one excluded class were mostly from “No.15: truck” class to “No:6 car” class.

**Figure 31:**
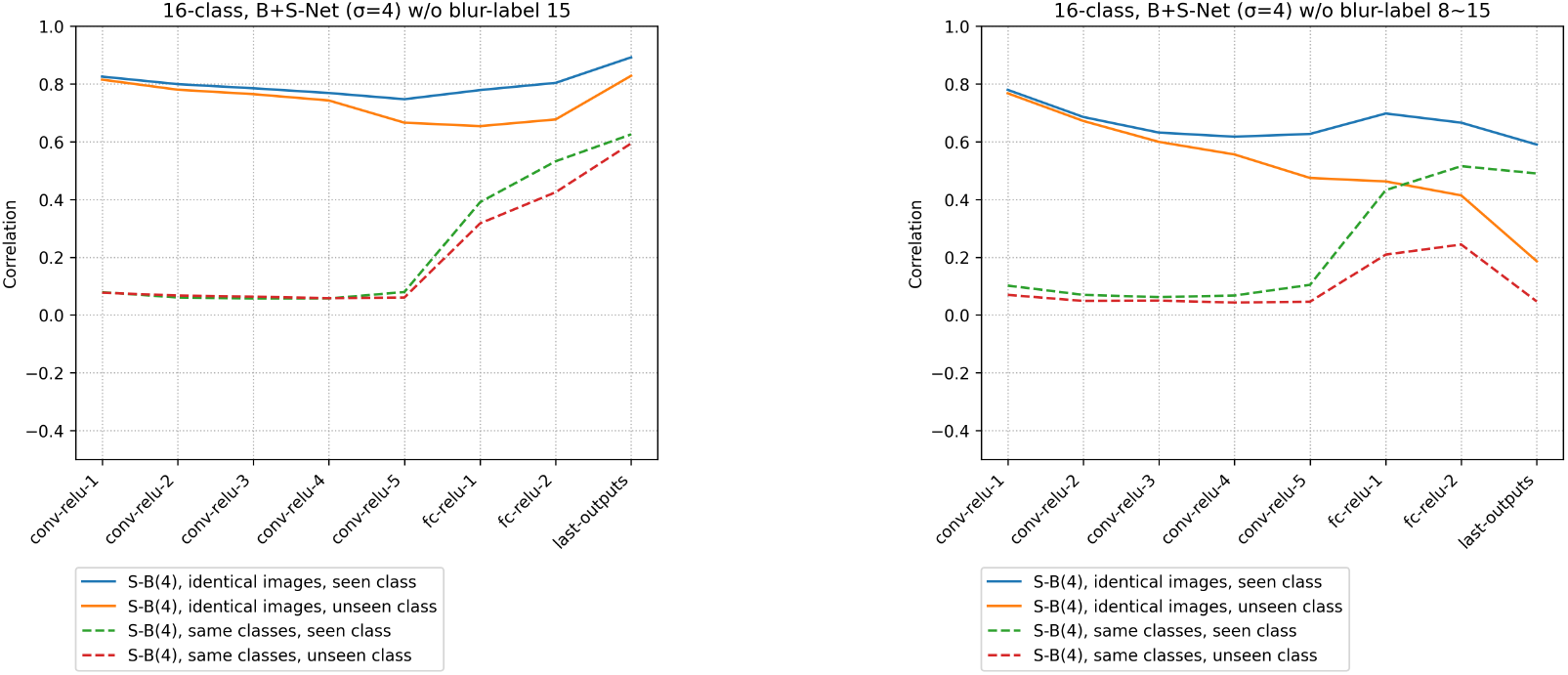
Mean activity correlation of the middle layer units for the sharp (S) and blurred (B) images of the B+S-Net trained without some blur labels. (left) model trained without a blurred label (No. 8), (right) model trained without some blurred labels (No. 8-15). identical images, seen class: mean activity correlation of identical images (learned class). identical images, unseen class: mean activity correlation of identical images (unlearned class). same classes, seen class: mean activity correlation with non-identical images of the same class (learned class). same classes, unseen class: mean activity correlation with non-identical images of the same class (unlearned class).

**Figure 32:**
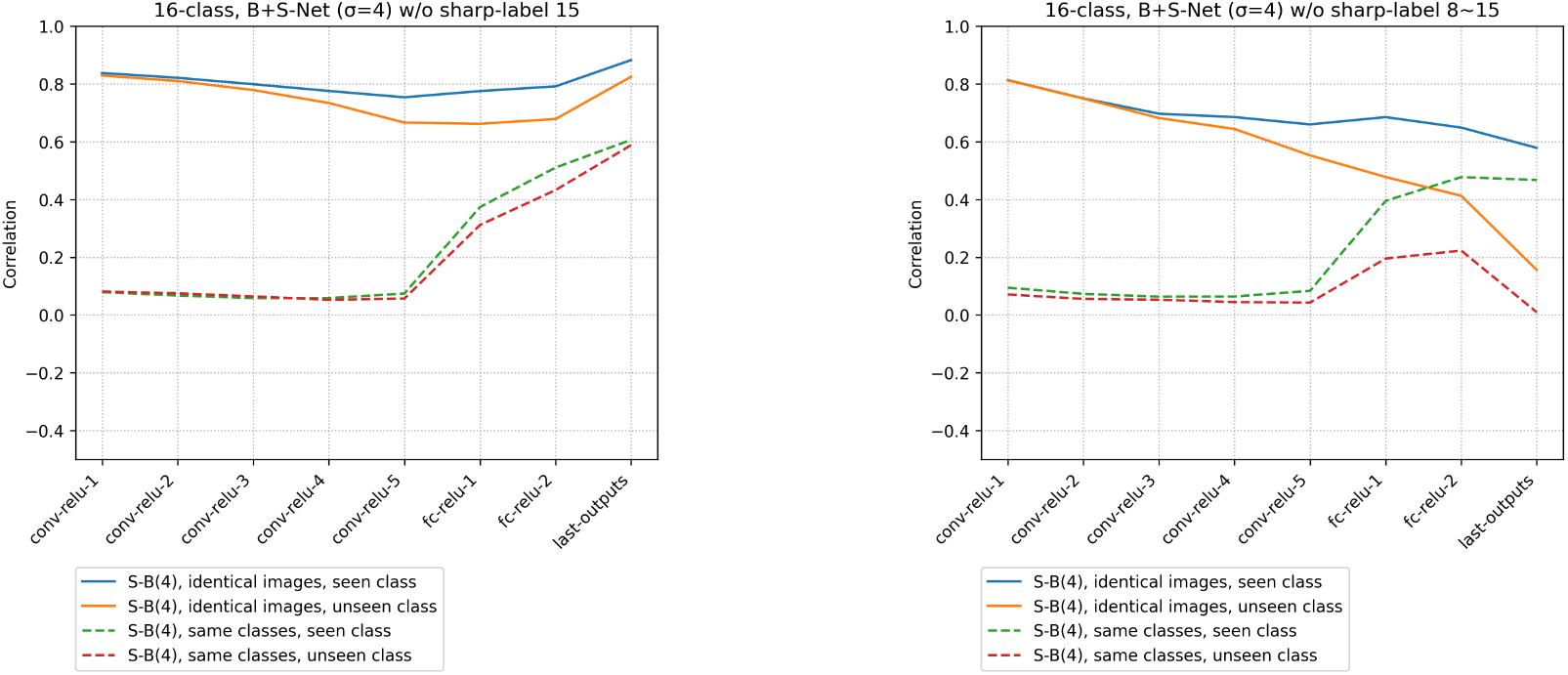
Mean activity correlation of the middle layer units for the sharp (S) and blurred (B) images of the B+S-Net trained without some sharp labels. (left) model trained without a sharp label (No. 8), (right) model trained without some sharp labels (No. 8-15). identical images, seen class: mean activity correlation of identical images (learned class). identical images, unseen class: mean activity correlation of identical images (unlearned class). same classes, seen class: mean activity correlation with non-identical images of the same class (learned class). same classes, unseen class: mean activity correlation with non-identical images of the same class (unlearned class).

Overall, the zero-shot transfer analysis suggested that the shared representation acquired by the blur training can be reused, at least partially, to recognize an object class with unseen image type (either blur or sharp) during training. This further supported the view that common representations that are invariant to blurry and sharp image inputs are formed in the earlier stages of the visual processing by blur-training (Case 1 in Figure 20). In a similar way, humans might efficiently acquire blur robust representations to general object categories just by being exposed to blurry images of a limited number of objects.

### 4.3 Does multi-scale frequency integration occur?

In the previous section, we analyzed the internal representation of sharp and blurred images. Given that a sharp image contains low spatial frequencies and high spatial frequencies and other information, information of different (low and high) frequencies may be integrated when sharp and blurred features are processed commonly. In this section, we analyze whether the CNN models integrate the multi-scale frequency information internally.

#### 4.3.1 Visualization of the internal representation with t-SNE

In a study of spatial frequency information processing in the human visual cortex, Vaziri-Pashkam et al.[Vaziri-Pashkam et al., 2019] used multidimensional scaling (MDS)[Shepard, 1980]. They showed that the representation of low-frequency and high-frequency information is close to each other in the V1 field, and the same categories of information are represented close in the ventral occipitotemporal (VOT) region.

To investigate how the high- and low-frequency information is represented in the CNN, we applied the t-SNE[Laurens van der Maaten and Geoffrey, 2014](perplexity=30, iteration=1000) and visualized the activity in the middle layers, compressing it into two dimensions for each layer.

The visualization results (Figure 33) from 10 pairs of high-frequency (H) and low-frequency (L) images of the same image for each class show that the representation of high-frequency and low-frequency features appears to be closer in B+S-Net than in S-Net. However, there was no tendency for the same frequency bands to be similar in the lower layers and the same object categories to be similar in the higher layers, as reported in the spatial information processing analysis of brain activity [Vaziri-Pashkam et al., 2019]. This is consistent with our finding in section 3 that neither S-Net nor B+S-Net can recognize objects in high-pass images.

**Figure 33:**
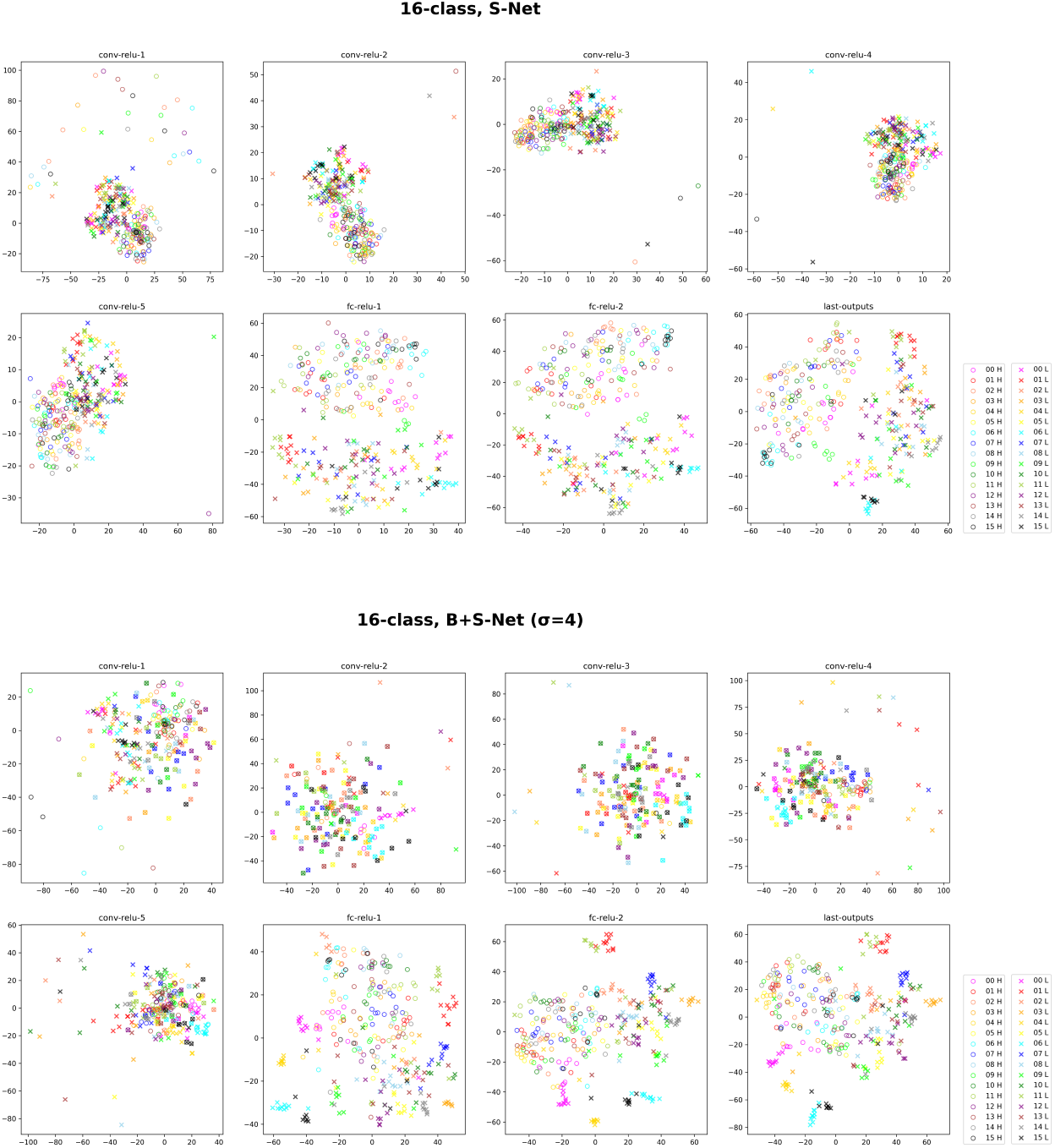
t-SNE visualization of the internal representation (high-frequency image (H) and low-frequency image (L))

#### 4.3.2 Activity correlation of the units of the intermediate layers

Next, we examined the average activity correlation (H-L correlation) of the middle layers between the high- and low-frequency images in the same way as in section 4.2.1.

As a result (Figure 34), the H-L correlation of the S-Net for the same image is low in the convolutional layers and slightly increased in the fully connected layers. It suggests that high-frequency and low-frequency features are processed separately in the convolutional layers. A part of the increase in the H-L correlation in the fully connected layers is insensitive to image types and thus possibly reflects a network bias. For the B+S-Net, there are weak positive H-L correlations for the same images in the convolutional layers.

**Figure 34:**
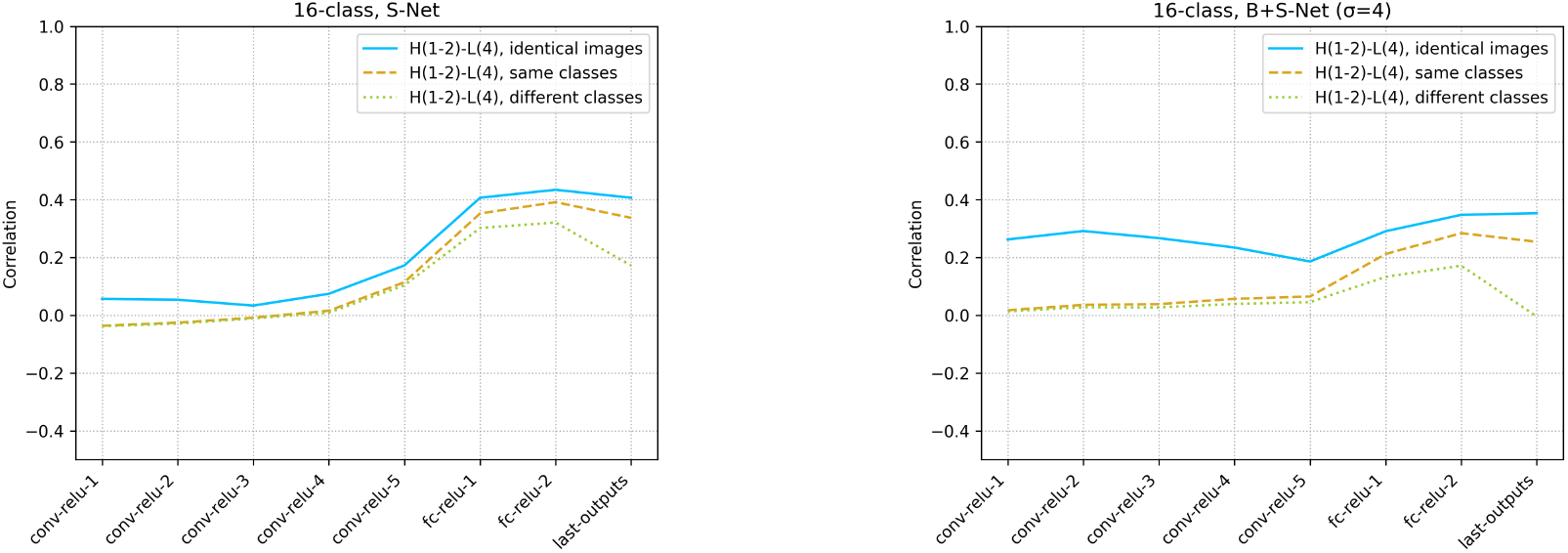
16-class-AlexNet average activity correlation of the middle layers units for high-frequency (H) and low-frequency (L) images

We also analyzed the H-L correlation of VOneNet, which has multi-scale Gabor filters in the first layer. The H-L correlation for the same images in the convolutional layers is low, and there is no clear evidence of integration of low and high frequency information, even after B+S training (Figure 35).

**Figure 35:**
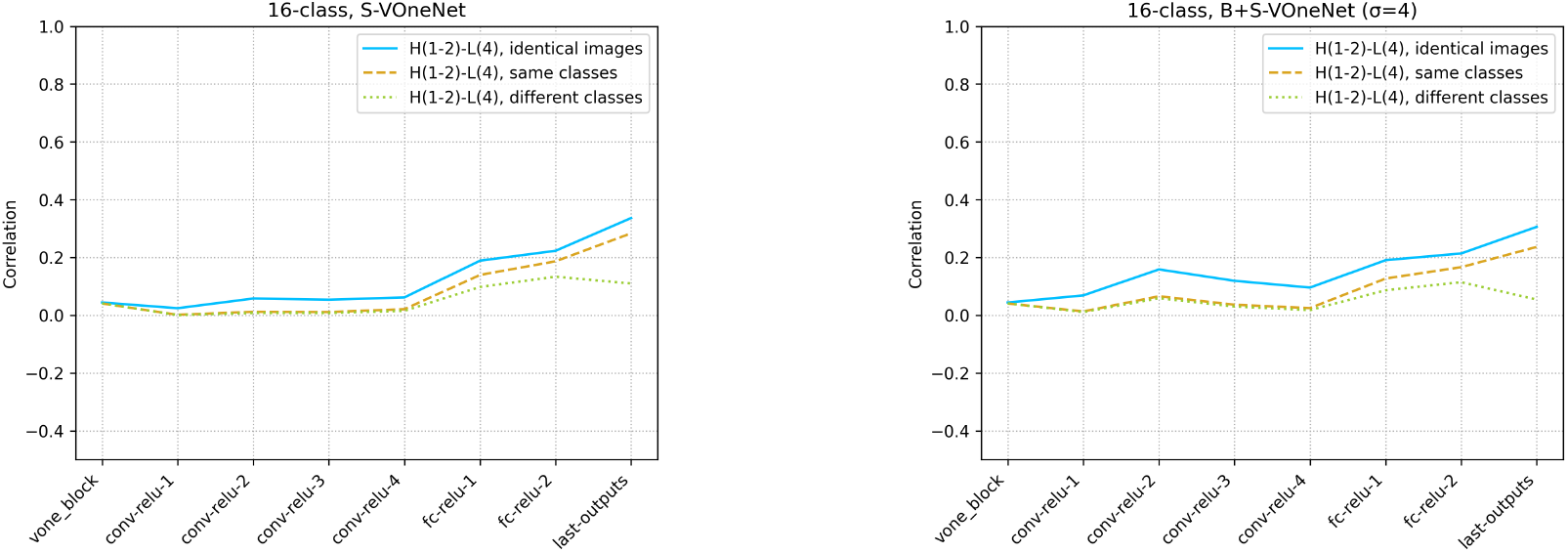
Average activity correlation of the middle layers units for high-frequency (H) and low-frequency (L) images of VOneNet

#### 4.3.3 RDM with band-pass image

The last analysis examined cross-frequency interactions in the CNN using only low-pass and high-pass images. To analyze cross-frequency interactions more in detail, we investigated how the information in each frequency subband is represented in the CNN by representation similarity analysis (RSA)[Kriegeskorte et al., 2008], computing the representational dissimilarity matrix (RDM)[Nili et al., 2014] with a range of band-pass images.

First, we created RDMs by activating a CNN with a range of band-pass images created from the same image and correlating the activity of the intermediate layers across subbands (Figure 36 upper). This allowed us to analyze the relative representation of each band-pass information in the intermediate layers.

**Figure 36:**
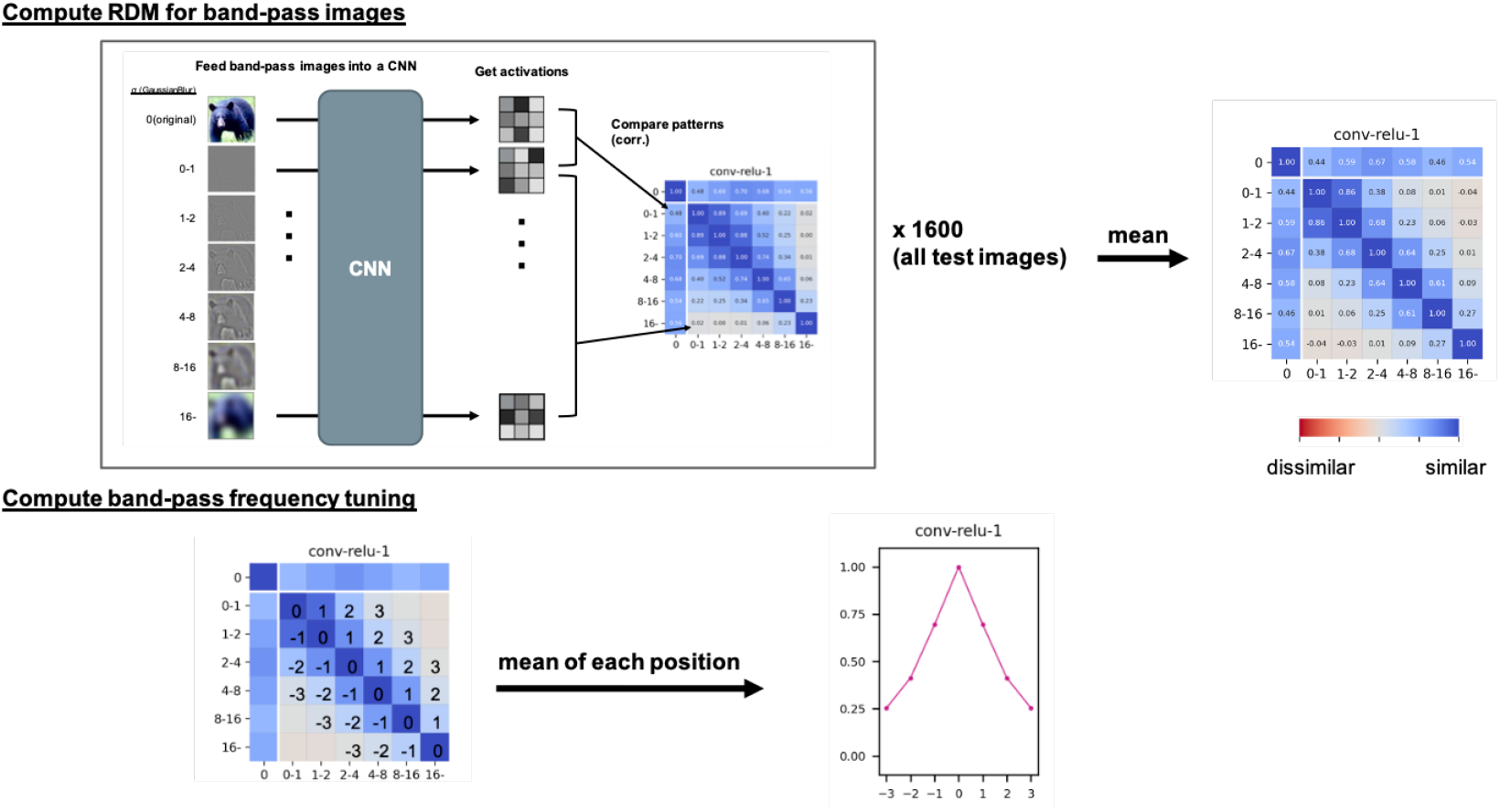
We compute RDM to revealbandpass frequency tuning of each layer. Each cell of the matrix represents the correlation of unit activation between different subbands. The original broadband image is indexed as 0 (leftmost column and the top row).

In order to visualize the band-pass frequency tuning of each layer, we computed the cross-subband correlation values as a function of relative frequency averaged over subbands of different center frequencies (Figure 36 bottom).

Figure 37 shows the RDM of S-Net and B+S-Net, and Figure 38 shows the frequency tunings, for 16-class AlexNets. First, the peak of correlation between the original broadband images and the band-pass images is shifted to lower frequencies for B+S-Net than for S-Net in both convolutional and fully connected layers. Second, the correlation between different subbands increases, and frequency tuning becomes broader, in the deeper layers. Third, as shown in Figure 38, the frequency tuning in the convolution layers is broader for B+S-Net than for S-Net.

**Figure 37:**
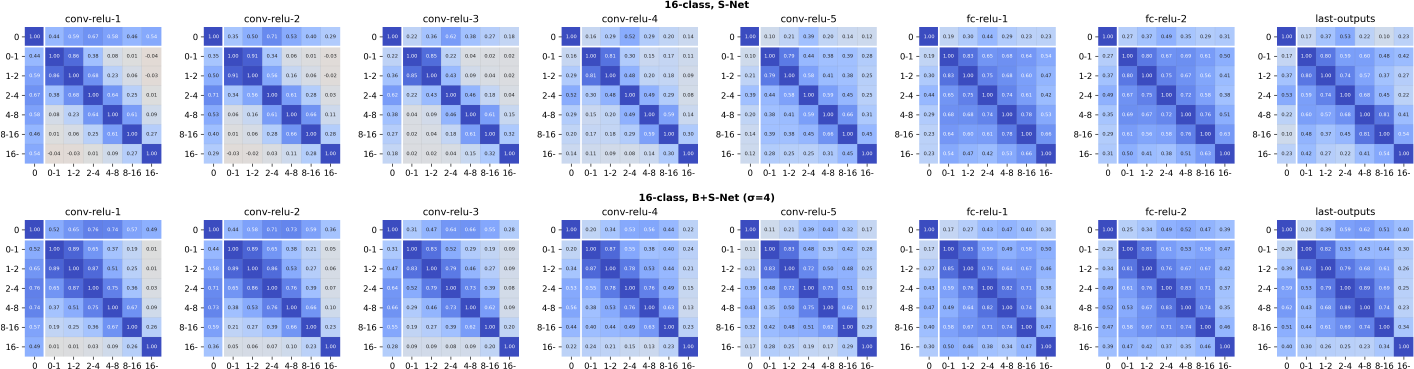
RDM (16-class-AlexNet)

**Figure 38:**
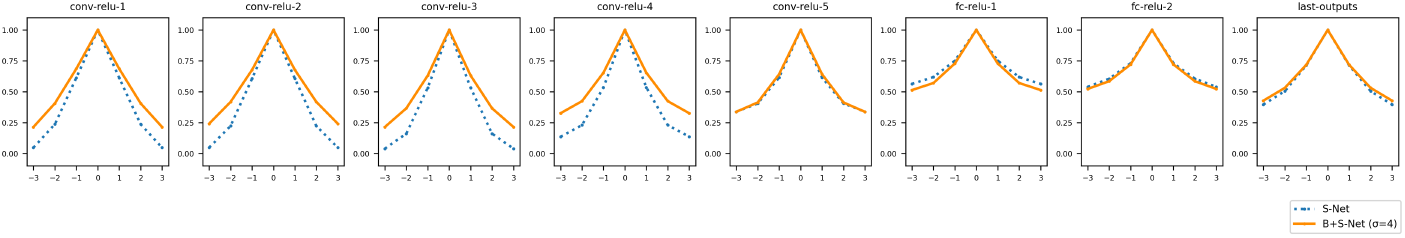
Band-pass frequency tuning (16-class-AlexNet)

These results suggest that subband information may be gradually integrated in the network, and that blur training assists this integration. However, for 1000-class-AlexNet, we did not find clear changes in frequency tuning among layers or between training types (Figure 39). In this case, CNN does not acquire frequency-invariant representations even when trained with a mixture of sharp and blur images.

**Figure 39:**
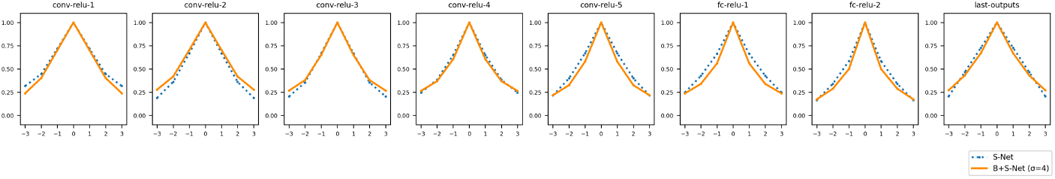
Band-pass frequency tuning (1000-class-AlexNet)

### 4.4 Summary and Discussion of Section 4

In this section, we analyzed how B+S-Net, which performed similarly to humans in a low spatial frequency image ficlassi cation task, processed sharp and blurred images.

The activity correlation between sharp and blurred images increased in B+S-Net. The results suggest that B+S-Net extracts more common features from sharp and blurred images than S-Net.

The results of zero-shot transfer learning support this view. While the generalization accuracy is not very high, the confusion matrix and the internal activity correlation suggest that B+S training produces a certain degree of common representation between blur and sharp features, which can be used even for unlearned categories.

These results suggest that B+S-Net recognizes sharp or blurred images using common representations, rather than using separate representations. The results also suggest that it is not only linear low-pass filtering in the first layer, but also a series of non-linear processing in the subsequent layers, that produces the common representations. In this aspect, we may be able to get useful computational insights into human processing from the analysis of B+S-Net.

Whereas we found B+S training facilitates the development of common processing for sharp (broadband) and blurred (low-pass) images, we found little evidence for B+S training facilitating the development of common processing for low-pass and high-pass images, nor integration of subband information. These results suggest that the frequency processing by B+S-Net is critically different from that by the human visual cortex. How can we make the frequency processing closer to that of the human visual system? Several machine learning techniques including data augmentation and contrastive learning may be used to force the network to integrate subband information. Note, however, that as a tool to understand human visual computation, it is important that the model training is natural and plausible for the development of human visual system, like blur training.

## 5 Conclusion

In this study, we investigated the effect of experiencing blurred images on forming a robust visual system to the environment as one of the factors for constructing an image-computable model of the human visual system.

To this end, we compared the recognition performance of CNN models trained with a mixture of blurred images using several different strategies (Blur-training).

The results show that the B+S-Net trained with a mixture of sharp and blurred images is the most tolerant of a range of blur and most human-like. In addition, the training in the order from blurred to sharp images, as claimed in the previous study [Vogelsang et al., 2018], was not very beneficial.

Other evaluations of the model’s performance with test stimuli showed that Blur-training did not improve the recognition of global spatial shape information, or only slightly.

B+S-Net extracts common features between sharp and blurred images. It however dos not show integration of multiscale (high and low frequency) frequency information, unlike in the human visual cortex.

In conclusion, training with blurred images provides performance and internal representation comparable to that of humans in recognising low spatial frequency images. It does narrow, but only slightly, the gap with the human visual system in terms of global shape information processing and multi-scale frequency information integration.

## Footnotes

1 The equation is implemented in OpenCV’s GaussianBlur function that we used to apply Gaussian filtering. In this function, the kernel size was adaptively determined from the size of sigma

2 texture-shape cue conflict image: taken from the GitHub page of [Geirhos et al., 2019](https://github.com/rgeirhos/texture-vs-shape/tree/master/stimuli/style-transfer-preprocessed-512, reference date: 2021/07/26)

3 The trained model was used directly in the analysis. Fine-tuning may yield different results.

## Notes

### Competing Interest Statement

The authors have declared no competing interest.

